# Multi-step genomics on single cells and live cultures in sub-nanoliter capsules

**DOI:** 10.1101/2025.03.14.642839

**Authors:** Ignas Mazelis, Haoxiang Sun, Arpita Kulkarni, Theresa Torre, Allon M Klein

## Abstract

Single-cell sequencing methods uncover natural and induced variation between cells. Many functional genomic methods, however, require multiple steps that cannot yet be scaled to high throughput, including assays on living cells. Here we develop capsules with amphiphilic gel envelopes (CAGEs), which selectively retain cells and large analytes while being freely accessible to media, enzymes and reagents. Capsules enable high-throughput multi-step assays combining live-cell culture with genome-wide readouts. We establish methods for barcoding CAGE DNA libraries, and apply them to measure persistence of gene expression programs in cells by capturing the transcriptomes of tens of thousands of expanding clones in CAGEs. The compatibility of CAGEs with diverse enzymatic reactions will facilitate the expansion of the current repertoire of single-cell, high-throughput measurements and extension to live-cell assays.

## Main Text

Single cell genomics encompasses a set of methods whereby hundreds to millions of cells are individually subject to highly-multiplexed assays including DNA sequencing, chromatin accessibility or modification, RNA sequencing, or combinations thereof (*1*, *2*). These methods enable unbiased, systematic discovery of cellular phenotypes and their dynamics, including immune cell responses, transitional states in tissue development, and changes in disease or upon perturbation. However, integrating these tools with each other or with functional cell screens faces limitations because the amount of material that can be obtained from each single cell is small and often requires reaction conditions incompatible with each other (*1*). Balancing sensitivity, scale, and genome-wide breadth with live culture capabilities imposes trade-offs between assay type, performance, and scalability.

Single cell sequencing methods are currently implemented via three technological modalities: in wells (e.g. SMART-Seq3) (*3*, *4*), in micro-compartments (droplets or nano-wells) (*5–7*), or by in-cell barcoding (ICB) of cross-linked cells (sciRNA-Seq and SPLiT-Seq) (*8*, *9*). These methods each offer different trade-offs between sensitivity and number of cells that can be analyzed, and they suffer different limitations on complexity of assays that they enable. Well-based methods are typically the most sensitive, possibly because each cell is processed in multiple steps in a manner resembling bulk methods. These methods also enable carrying out functional or live-imaging assays on living cells prior to genomic analysis. However, well-based methods scale poorly. Micro-compartment-based methods achieve increased throughput by physically isolating individual cells together with individual DNA-barcoded beads, enabling cell-specific barcoding of material from thousands of cells together within one sample. However, these methods are limited to a single reaction step once cells are compartmentalized. ICB methods scale to exponentially large numbers of cells, but require cell cross-linking, which leads to material loss and reduced measurement sensitivity (*8*, *9*). Neither droplets nor ICB methods allow parallelizing functional live-cell assays. Accordingly, approaches have been developed to pre-process cells prior to their analysis by these modalities, such as single cell ELISA (*10*, *11*) or DNA tagging of cell populations (*12*, *13*).

A technology that may unify advantages of these modalities are semi-permeable capsules (*14*, *15*). Capsules are sub-nanoliter spherical liquid compartments surrounded by a resilient shell. The chemical composition of the shell can be chosen to be selectively permeable to small molecules including tissue culture media, small proteins such as DNA polymerases and ligases, and oligonucleotides, while selectively retaining larger macromolecules like DNA or mRNA (**Fig. 1A**). Like tissue-culture wells, they have potential to allow culturing cells in isolation prior to analysis, with free exchange of media and nutrients. They also allow carrying out optimized molecular biology reactions on compartmentalized cells or analytes by enabling the repeated exchange of buffers and reagents. Like droplets, capsules can be produced at scale and their resilient shell allows sorting to enrich cells or colonies with desired features (*15*). These properties expand the range of genomic assays that are feasible for single cells and live cell cultures and present a versatile technological platform that may allow parallelizing assays that until now have been difficult and costly to carry out at high throughput (**Fig. 1B**). Here we report the development of capsules with appropriate permeability and stability for genomic assays, implement a split-pool barcoding-based approach to generate single-capsule genomic and transcriptomic libraries, and apply these developments to perform a massively-parallel live-cell assay characterizing the diversity and persistence of transcriptome-wide expression fluctuations in clones. The utility of capsules for multi-step genomics is also shown in a co-submitted paper (Baronas et al, co-submission), exploring different applications.

**Fig. 1.**
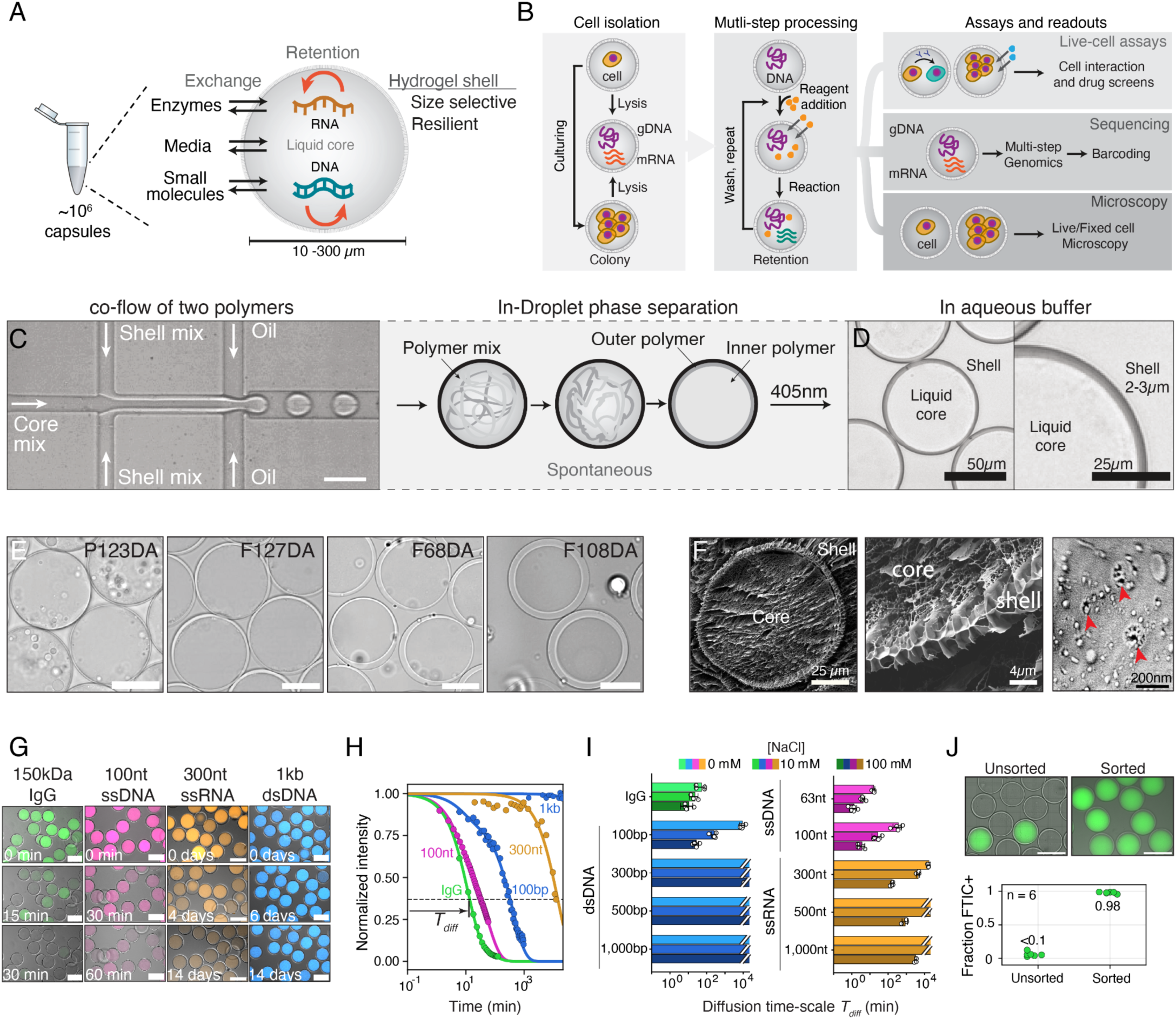
Concept, development, and characterization of pluronic diacrylate capsules. **(A)** Schematic of semi-permeable capsules and their relevant properties for high-throughput, single-cell profiling. **(B)** Schematic of potential massively-parallel, multi-step analyses in capsules including processing of single or cultured cells through iterative addition and removal of reagents, imaging, sorting, and sequencing. **(C)** Bright-field micrograph (left) and schematic (right) showing generation of CAGEs (compartments with amphiphilic gel envelopes) using a droplet generator co-flowing an amphiphilic functionalized shell polymer and a hydrophilic core polymer that undergo phase-separation upon droplet formation, and are subsequently converted to CAGEs by 405nm-dependent cross-linking of the shell polymer. Scale bar=100µm. **(D)** Bright-field micrographs of 8% (w/v):1% (w/v) F127DA:PPPDA shell, 13% (w/v) dextran core CAGEs in DPBS. **(E)** Bright-field micrographs demonstrating that multiple amphiphilic PEG copolymers form capsules. Scale bar=50µm. **(F)** Cryo-SEM images of freeze-fractured capsules with magnification of pores in the pluronic diacrylate shell membrane. Red arrows point to surface pores. **(G)** Illustrative composite micrographs of confocal and bright-field imaging of capsules from analyte diffusion time-series. **(H)** Intensity time-series quantifying diffusion half-times through the CAGE shell (composition as in **D**) for analytes of different sizes and in the presence of different salt concentrations. **(I)** Diffusion half-times showing a sharp size-dependent cut-off in CAGE permeability. Plot represents mean data from ≥3 independent experiments with SD as error bars. Full data in **Table S1**. **(J)** Flow cytometry–based enrichment of CAGEs. Composite fluorescence and bright-field micrographs of a sample initially containing <10% labeled capsules (top-left) and sample after sorting (top-right), yielding ∼98% FITC-positive CAGEs (bottom). Scale bar = 100 µm.

### Developing capsules with amphiphilic gel envelopes (CAGEs)

Capsules are generated by forming concentric layers of core and shell components (*16*), with one approach using liquid-liquid phase-separation (LLPS) between two polymers inside microfluidic droplets(*14*, *15*). The outer polymer can then be cross-linked to form a stable shell. For biological applications, several studies have reported production of capsules consisting of a poly-ethylene glycol diacrylate (PEGDA) shell (*14*, *15*). However, PEGDA fails to form stable shells under changes in buffer pH, salinity, and concentration, thereby limiting the ability to control capsule permeability (*17*, *18*) (recapitulated in **Fig. S1A**). Amphiphilic molecules should more reliably wet the water-oil interface of a droplet than PEG, so we speculated that such molecules would be more suitable for producing stable concentric shells through spontaneous LLPS. We tested the addition of Pluronic F127, a biocompatible and FDA approved amphiphilic block-copolymer, and found that it significantly improves the production of PEGDA capsules in different conditions (**Fig. S1B, C**). By functionalizing Pluronic F127 with diacrylate (F127DA), it can serve as a shell building block on its own (**Fig. 1C**). The production of pluronic diacrylate (PDA) capsules occurs through LLPS between PDA and dextran inside simple microfluidic droplets. Once formed, the PDA is cross-linked into a physical gel by use of a photoinitiator and brief exposure to 405nm light (**Fig. 1C**). The resulting PDA capsules are optically clear (**Fig. 1D**), can be reliably formed at scale and are tunable by varying polymer concentrations (**Fig. S1D**). Additionally, the strategy is generalizable: capsule production is possible with other amphiphilic copolymers of different lengths, including other Pluronics (P123DA, F68DA and F108DA) (**Fig. 1E**), enabling the formation of capsules with varying shell properties. We refer to these capsules as CAGEs: capsules with amphiphilic gel envelopes.

We next optimized the selective permeability of CAGEs. We focused on using F127DA as Pluronic F127 undergoes micellization (radius 10-20 nm (*19*)), and F127 hydrogels have been shown to form a packed micellar array (*20*). Genomic assays utilize small enzymes and oligonucleotide primers of up to ∼100 nt in length which are estimated to have a gyration radius <10 nm (*21*)), while the dsDNA generated during library preparation are larger (300bp or longer, persistence length >50 nm for dsDNA (*22*)). Indeed, Cryo-SEM imaging shows that the F127DA shell has a foam-like structure, with evidence of surface pores of typical diameter of ∼10-50 nm (**Fig. 1F**). We generated CAGEs with a shell composed of F127DA and other copolymers, and measured mean life time (*T*_Diff_) associated with the diffusion of fluorescently-labeled proteins, DNA and RNA through the CAGEs shell (**Fig. 1G, S1E-G**). We identified a starting polymer composition for capsule formation for which dsDNA of length ≥300 bp showed no diffusion through the CAGE shell (diffusion lifetime *T*_Diff_ >14 days) in any buffer tested, while immunoglobulins and 100 nt ssDNA were able to diffuse into the CAGEs (*T*_Diff_ ∼5 min) (**Fig. 1H,I, Table S1**). For those molecules which exhibited diffusion, we further observed slower diffusion with decreasing salt concentration **(Fig. 1I**), and with increasing temperature (**Fig. S1I**). This makes PDA CAGEs particularly suitable for high temperature reactions, such as PCR (**Fig. S1E**), where retention of molecules at high temperature is desirable. The observed ramp in diffusion with increased salt concentration also suggested a strategy for modifying buffers for washing, storing or loading CAGEs. Further, F127DA offers a stable base for capsule formation, meaning that diffusion rates can be easily tuned by altering polymer composition (**Fig. S1H**).

Overall, the resilient shell of CAGEs allows them to be manipulated through multiple steps of analysis. CAGEs can be handled with common lab equipment, can be analyzed by flow cytometry, and can be sorted through a commercial FACS instrument to selectively fractionate them based on fluorescence (**Fig. 1J**). Taken together, these results demonstrated that CAGEs offer a physically robust platform, which possess a tunable permeability threshold enabling diffusion of primer-scale ssDNA and proteins while retaining large macromolecules, making them suitable for multi-step molecular biology reactions and thus high-throughput genomics.

### Barcoding DNA in capsules

We reasoned that CAGEs can enable massively-parallel analyses by first compartmentalizing live cells or macromolecules and subsequently performing sequential enzymatic reactions interposed with wash steps between each step to introduce new reaction conditions. This approach would require labeling the analytes in each compartment through DNA barcoding that later identifies their capsule-of-origin by sequencing.

Using CAGEs, we developed a flexible split-and-pool barcoding (*8*, *9*) approach that allows indexing of double-stranded DNA molecules—the final form of all major analytical libraries analyzed by sequencing (**Fig. 2A**). The approach first amplifies the DNA derived from different assay reactions, such as RT or Tagmentation, to overcome inefficiencies in subsequent barcoding. This amplification is carried out by PCR with one of the two primers contains a deoxyuridine (dU) residue proximal to its 3′ end, which can be efficiently cleaved by dU-excising enzyme to remove a PCR handle and generate a 4-nt 3′ overhang and a 5′ phosphate used subsequent barcoding (**Fig. 2B, S2A**). This process can be iteratively repeated after each barcode addition to remove linker DNA sequences and to generate a 5′-P ligation site for the next round. In this work, CAGEs are subject to two rounds of split-pool ligation and one round of PCR DNA barcoding (schematic in **Fig. 2A**) after which the prepared libraries are readily released by dissolving CAGEs (**Fig. S2B,C**). Additional barcodes can be added during initial library construction (via PCR, RT, or Tagmentation) as well as indexing after release from CAGEs, scaling the number of barcodes. The number of capsules that can be profiled scales exponentially with the number of reaction steps (**Fig. S2D, E**). We refer to this method for indexing DNA libraries in capsules as inC-seq.

**Fig. 2.**
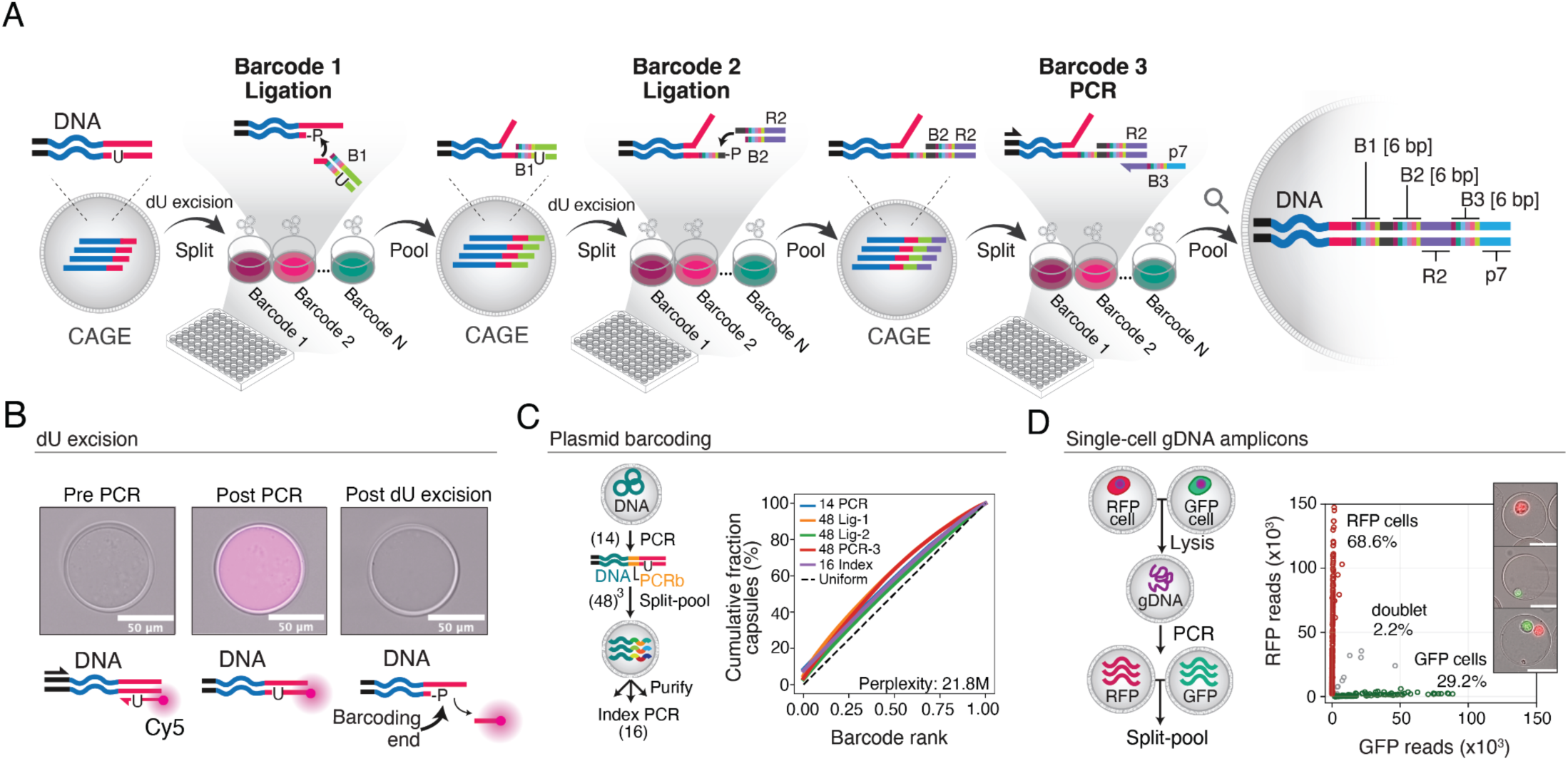
Split-and-pool-based barcoding of CAGEs for high-throughput sequencing (inC-Seq) **(A)** Schematic for split-and-pool barcoding of DNA libraries in capsules, showing an example of three rounds of barcoding through ligation and PCR. Initial libraries are generated with a uracil residue in the common library end-sequence (red), which is cleaved by a uracil-specific excision reagent to generate a sticky end with a 5’-phosphate. The sticky ends are ligated to an array of barcoding dsDNA oligos that contain a well-specific barcode. The process is repeated to introduce a second barcode, and a third barcode is introduced through a PCR primer. **(B)** Demonstration of efficient sequential enzymatic processing of DNA in CAGEs, realizing step (1) from panel (A). An initial unlabeled library (left panel) is amplified by incubation of capsules in a PCR reaction mix, with a PCR primer containing a uracil and a Cy5 label at its 3′ terminus. Following PCR and washing (middle panel), the CAGEs show retention of Cy5 fluorescence that now is attached to the DNA in the capsule. Uracil excision and washing cleaves the labeled primer generating an overhang and losing Cy5 fluorescence (right panel). **(C)** Representation of barcodes in the DNA amplicon sequencing data after a 5-step barcoding approach: 14 PCR barcodes (PCRb) were added during amplification, followed by 3-steps of 48 split-pool barcode addition and 16 indices following capsule solubilization. **(D)** Effective library partitioning by capsules with low cross-contamination seen by inC-Seq analysis of single-cell gDNA transgenes loci encoding GFP or RFP sequences. The plot shows the number of reads from mapping to GFP or RFP, with each point corresponding to a unique capsule barcode. Inset displays examples of CAGEs with cells prior to lysis. Scale bar=50µm

We tested five-step inC-seq barcoding on CAGEs. We encapsulated plasmid DNA and then barcoded the capsules with 14×48×48×48×16=24.7 ×10^6^ potential barcodes. Applying this protocol to 350 µL of packed CAGEs, we obtained 355,418 capsule barcodes (1µL corresponding to ∼1000 capsules) with each of the possible barcodes elements uniformly represented across all five round of barcode addition, yielding an an effective theoretical barcoding space (Shannon diversity index) of 21.8 million barcodes (**Fig. 2C**). We next tested inC-seq in combination with digital PCR on single-cell derived material by isolating individual cells from a mixture of cells carrying transgenes encoding red or green fluorescent proteins (RFP or GFP) into CAGEs and then amplified these loci from genomic DNA (gDNA) (**Fig. 2D**). The DNA libraries produced from cells demonstrated that the cells and genomic material within each CAGE were effectively compartmentalized through the entire multi-step barcoding protocol, with 99±5% (median±SD) of reads from each capsule mapping uniquely only to single GFP or RFP sequences (**Fig. 2D**).

### Single-cell genomic assays in CAGEs

DNA barcoding in capsules enables diverse analytical approaches to study DNA and RNA. We implemented single capsule transcriptomics (inC-RNA-seq) and, separately, a single capsule assay for transposase-accessible chromatin (*23*) (inC-ATAC-seq) (**Fig. 3A,B; S2E**). We used a mixture of human and mouse cells to confirm the effective isolation of cellular material in each CAGE and a low rate of cell doublets (**Fig. 3C,D**). For inC-ATAC-seq, we observed a characteristic nucleosome peak in the fragment size distribution, and a strong enrichment of fragments at transcription start sites (17-fold enrichment) with low mitochondrial reads (<3%) (**Fig. S3A)**.

**Fig. 3.**
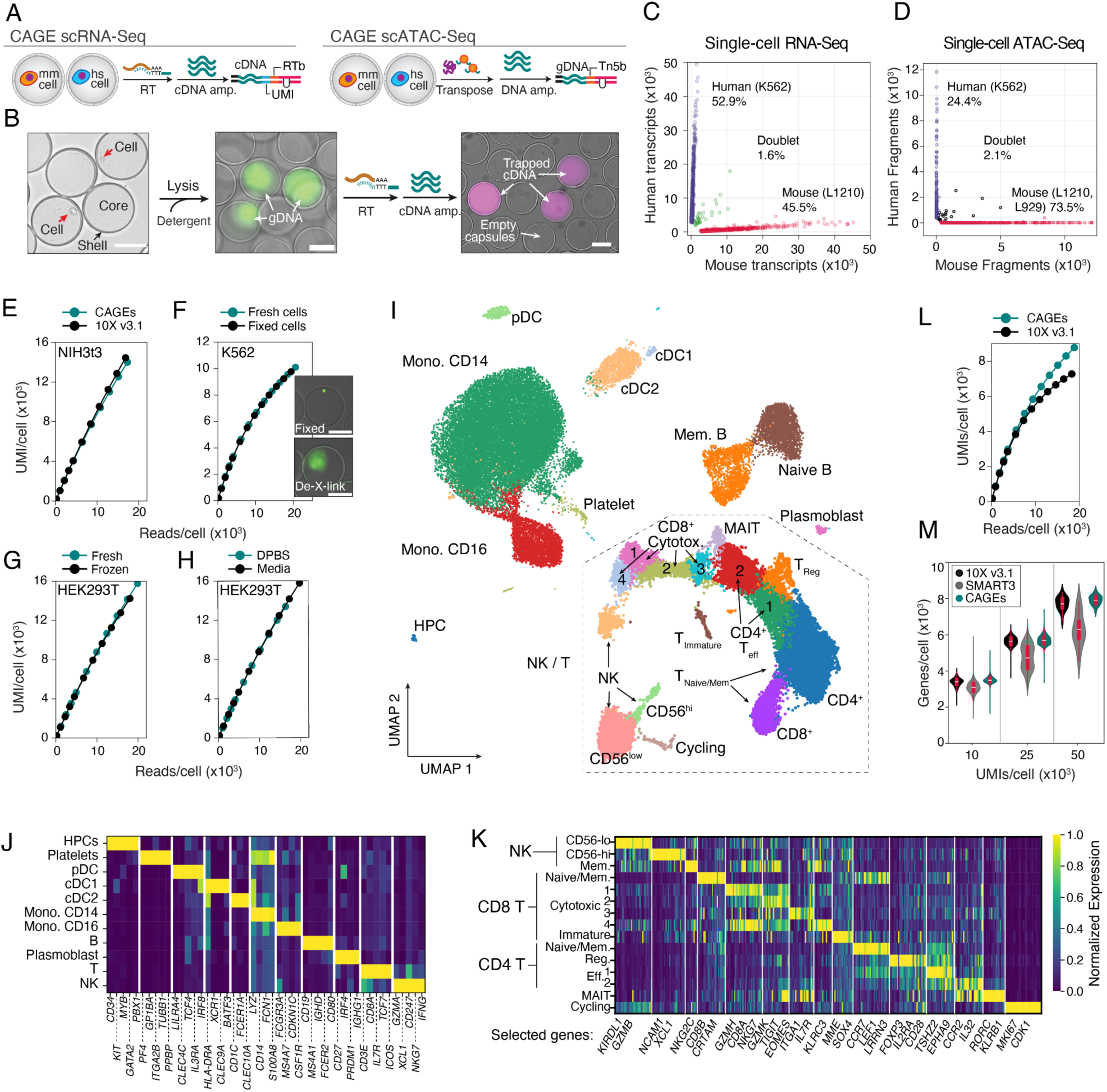
Capsules enable single-cell genomic assays with multi-step processing. **(A)** Schematic of inC-RNA-seq and inC-ATAC-seq library preparation modalities. Additional barcodes and sequences are added during transcript (RTb and UMI) and accessible gDNA region (Tn5b) capture via RT and Tagmentation steps. Captured and amplified dU bearing molecules are subjected to split-pool barcoding and sequencing. **(B)** Micrographs showing cells at progressive stages of single cell RNA-seq library preparation in CAGEs, from initial cell encapsulation through lysis, to generation of amplified complementary DNA (cDNA) libraries prior to in CAGE barcoding. gDNA is stained with SYBR Safe dye; cDNA stained with Cy5 fluorophores conjugated to the PCR amplification primers. **(C-D)** Plots showing the number of reads corresponding to each CAGE mapping to human (y-axis) and mouse (x-axis) RNA-seq (left) or ATAC-Seq (right) inC-seq libraries. The doublet rate is <5% as expected for Poisson loading of ∼0.1. **(E)** UMI/read plots following downsampling reads from CAGE scRNA-seq libraries (teal) in comparison to publicly available 10× Genomics NextGem v3.1 data (black). **(F, G, H)** UMI/reads plots as in (E) for libraries generated from fresh or PFA fixed K562 cells **(F),** fresh or frozen HEK293T cells **(G)** or capsules made using DPBS or media **(H)**. **(I)** A two-dimensional UMAP embedding of inC-RNA-Seq data from PBMCs, (**J**) heatmaps of gene expression showing coverage of low and high-abundance cell type-specific genes and **(K)** high population structure of partitioning subsets of T and NK cells. **(L)** UMI/reads plots as in (f), now generated for PBMC data and comparing inC-Seq to data from 10X Genomics GEMCode. **(M)** Comparison of number of genes detected per cell for inC-Seq data (teal) to publicly available 10X Genomics NextGem v3.1 (black) and SMARTseq3xpress data (*4*) (gray) PBMC datasets at three different levels of UMI/cell.

We then focused on optimizing methods for high sensitivity single-cell inC-RNA-seq, using the SMART-seq3 approach as a starting point (*3*). The flexibility of CAGEs allowed us to optimize sequential steps of lysis, reverse transcription and pre-amplification in cell lines (HEK293T and NIH3t3), to define a protocol that captures as many transcripts and genes as a commercial droplet-microfluidic system (10X Genomics, GEMCode v3.1) (**Fig. 3E, S3B,C**). We observed high reproducibility between independently prepared and barcoded capsules (**Fig. S3D**, log-correlation *R>*0.99) and between CAGEs and GEMCode (**Fig. S3E,** log-correlation *R*=0.95). Importantly, multi-step processing of CAGEs raises a question of whether smaller mRNA transcripts might be under-represented, but we find no apparent bias in the length of detected mRNA transcripts in capsules, in line with our diffusion assessment data (**Fig. S3F**). Further, we observed similarly high UMI-counts and reproducibility across replicates, as well as similarly low length-bias when performing inC-RNA-seq with: (i) PFA-fixed cells for which cross-linking was reverted in CAGEs prior to reverse transcription (**Fig. 3F, S3G**), (ii) material cryopreserved after cell lysis in CAGEs (**Fig. 3G, S3H**); and (iii) cells encapsulated while still in media—an inhibitor of RT—rather than after washing in PBS (**Fig. 3H, S3I**). These results indicate the versatility of CAGEs for complex assay design, and suggest their utility for profiling difficult-to-process samples, such as microbes (*14*, *15*), cells from marine organisms (*24*) or clinical PFA fixed samples.

To complete benchmarking inC-scRNA-seq, we profiled human peripheral blood mononuclear cells (PBMCs), which have been frequently analyzed by other methods. inC-RNA-seq of 43,665 PBMCs revealed the full landscape of PBMC states as seen by a UMAP representation (**Fig. 3I-K**), including rare circulating CD34+ progenitors, plasmoblasts and a clear partitioning of subsets of T cells and natural killer (NK) cells. Significantly, the method captured at least as many transcripts and genes as the commercial 10X Genomics GEMCode v3.1 and SMART-seq3 approach (*3*) (**Fig. 3L, M**) and considerably more transcripts than reported with other split-and-pool ICB methods (*9*, *25*). Overall, inC-seq approaches provide a versatile platform for high-throughput genome-wide measurements with the performance matching or exceeding the established transcriptomic profiling methods, and with the added flexibility of conducting multi-step reactions.

### Massively-parallel live-cell assays in CAGEs

Capsules have the potential to enable highly parallel functional assays in which cells are allowed to grow and/or interact prior to molecular profiling. To enable cell culturing in CAGEs, we modified our protocol to buffer against reactive oxygen species (ROS) formed during shell photopolymerization (*26*), which otherwise led to large-scale death of encapsulated cells (**Fig. S4A, B**). With this addition and no composition changes, cells captured in CAGEs are viable and clonally expand as demonstrated with several cell lines (L1210, K562, Jurkat, HEK293t, L929) (**Fig. 4A)**. CAGEs also support viable growth cells derived from primary mouse bone marrow-derived hematopoietic stem cells and human induced pluripotent stem cells (hiPSC), with latter forming cyst-like structures (**Fig. 4A**) and maintaining expression of *SOX2,* a marker of pluripotency **(Fig. S4C)**. Importantly, CAGEs can be produced at >2M/hour and thus enable high-throughput expansion of isolated colonies at scale (**Fig. 4B, S4D**), which can then be analyzed with the established inC-RNA-seq.

**Fig. 4.**
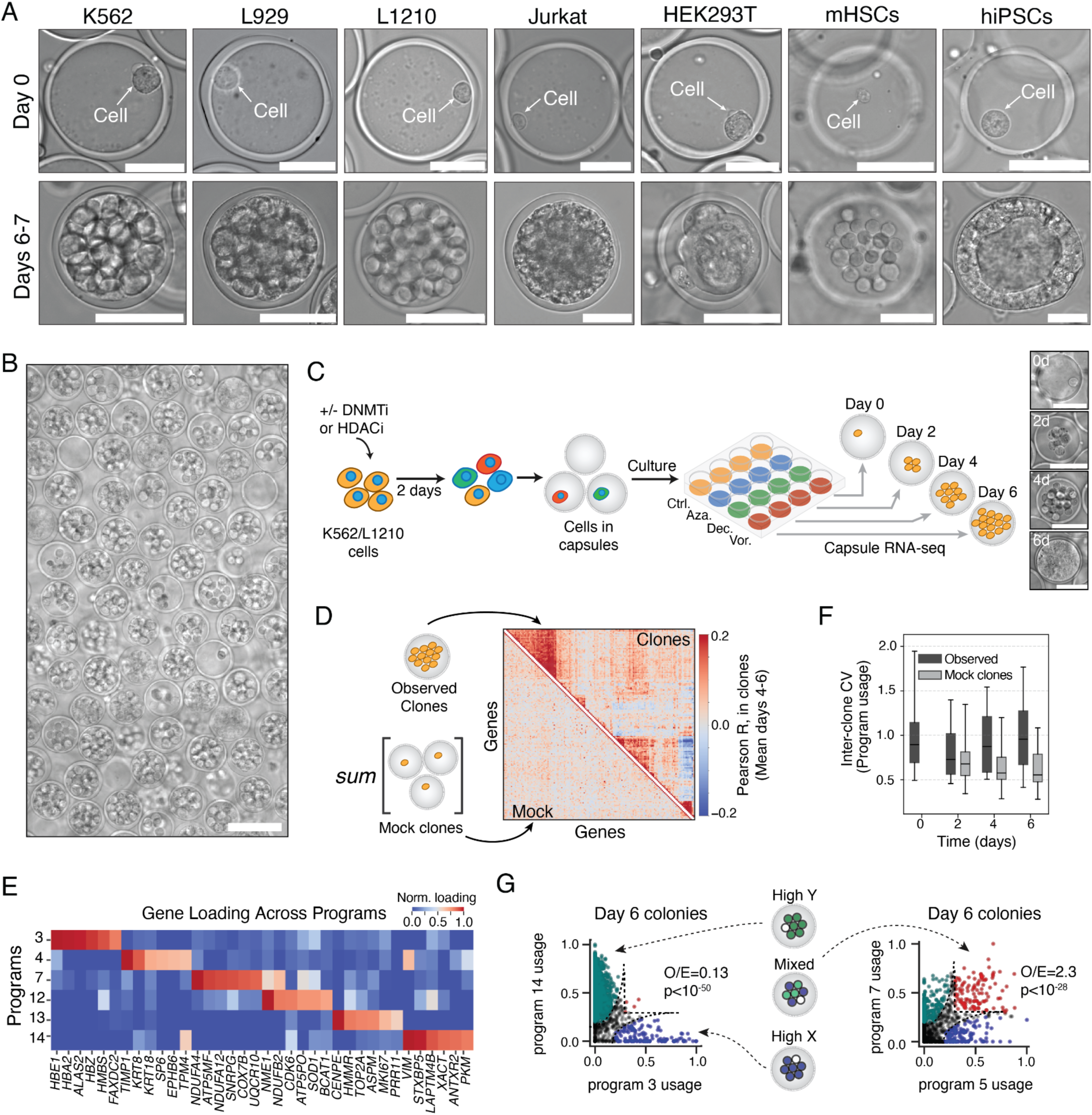
Live-cell clonal expansion in CAGEs reveals persistent gene expression programs. **(A)** Micrographs showing growth in CAGEs of clones from cultured cell lines, primary mouse bone marrow hematopoietic stem cells (mHSCs) and human induced pluripotent stem cells (hiPSCs). Scale bars = 50µm.**(B)** Micrograph of encapsulated colonies concurrently cultured. Scale bar = 100µm. The image shows high-density K562 (poly-clonal) colonies expanded in CAGEs. **(C)** Experimental schema for identifying persistent gene expression programs in cancer cell lines upon treatment with vehicle (Ctrl) or DNMT and HDAC inhibitors [5-azacytidine (Aza); decitibine (Dec), vorinostat (Vor)]. Micrographs show example capsules at different timepoints. **(D-G)** Evidence of persistent gene expression programs in K562 cells; see **Fig. S5** for L1210 cells. **(D)** Clustered inter-clonal gene-gene expression correlations in K562 cell control samples at days 4-6 show evidence of structured programs, which are absent when randomly combining cells sampled at day 0 into ‘mock’ clones. **(E),** Top genes contributing to gene expression programs identified by non-negative matrix factorization (NMF), identifying erythroid, cell cycle, vimentin- and keratin-associated programs. Full program loadings in **Table S2. (F)** Box plots of variation in 15 NMF-derived programs between control clones, showing persistent heterogeneity over time as compared to mock clones. **(G)** Some gene expression programs remain mutually exclusive after clonal expansion, as seen from low numbers of clones observed mixed NMF programs as compared to expectation (Observed/Expected = *f_XY_/f_X_f_Y_*; p-value from Fisher’s exact test).

We utilized this ability to grow cells in capsules in order to measure persistent clonal heterogeneity in gene expression. In several cancer types, cells occupy persistent epigenetic states, which are thought to underlie non-genetic heterogeneity in drug resistance (*27*, *28*). Drugs targeting the inheritance of epigenetic modifications (DNA methyltransferase (DNMT) inhibitors (*29*); and histone deacetylase (HDAC) inhibitors (*30*)) have been proposed as tools to activate tumor suppressor genes that are silenced, and they are used in combination with more traditional chemotherapies (*31–33*). Whether these drugs broadly reduce epigenetic persistence in gene expression, however, has not been explicitly tested. To measure the duration of epigenetic persistence, other studies have used RNA-seq to identify clonally-variable gene expression patterns – an approach so far carried out by culturing isolated cells in wells, with analysis restricted to a few dozen clones (*34*, *35*). Using CAGEs, one can parallelize this live-cell assay to carry out tens of thousands of parallel clonal growth experiments in parallel. We did so with cell clones derived from a mixture of human erythroleukemia (K562) and mouse lymphoblastoma (L1210) cells, and evaluated changes in clonal heterogeneity in the presence or absence of the DNMT inhibitors [decitabine (Dec) and 5-aza-cytidine (Aza)], and the HDAC inhibitor vorinostat (Vor) (**Figs. 4,5**).

**Fig. 5.**
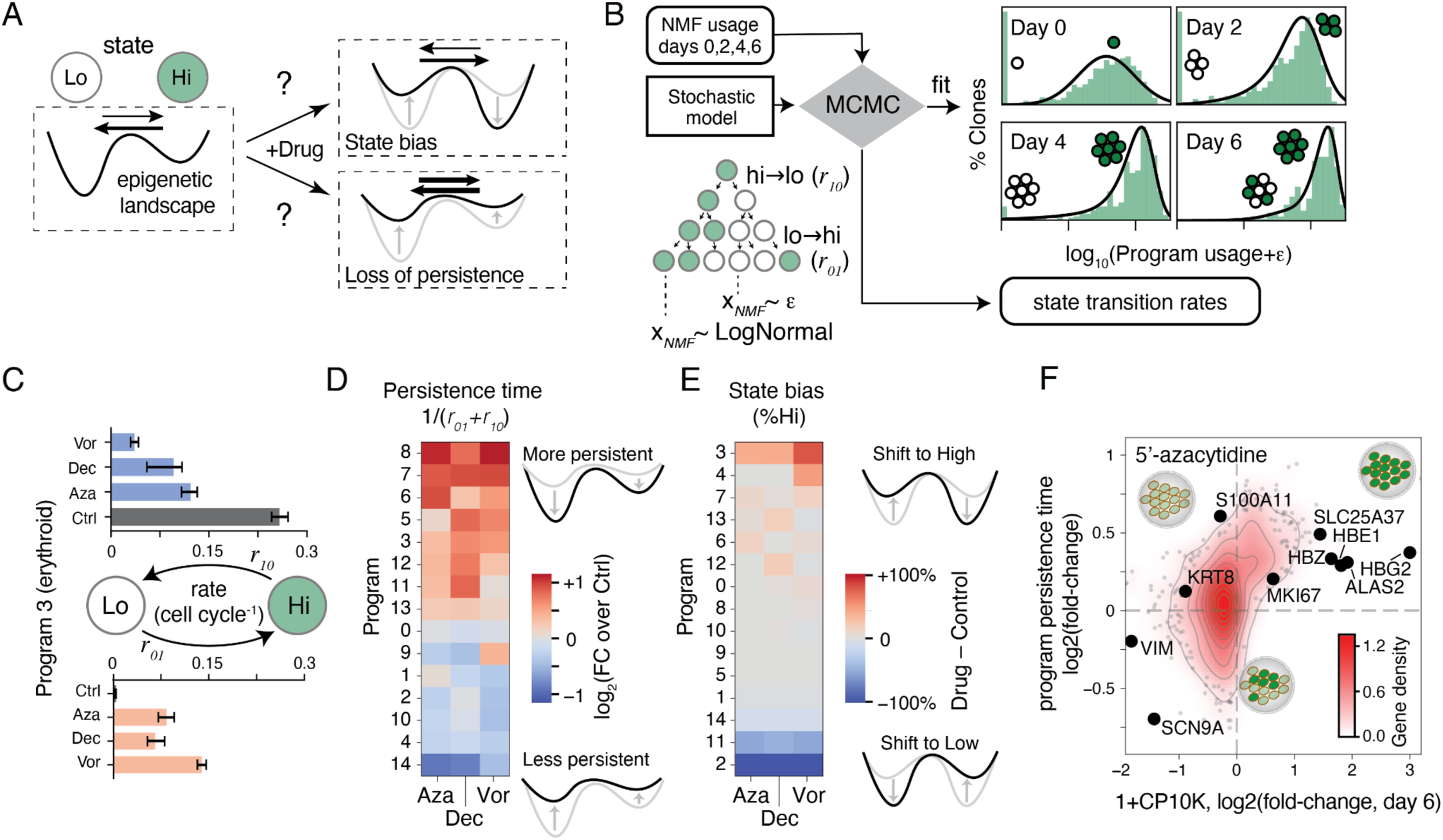
Altered clonal memory upon treatment with DNMT or HDAC inhibitors. **(A)** A schematic of possible effects of drug treatment on epigenetic memory. In untreated controls, cells occupy high and low gene expression states for each observed program. Inhibitors of epigenetic modifying enzymes may bias the relative stability of states (top scenario), and may also reduce persistence (shallower wells, bottom scenario). From inC-RNA-Seq on growing clones one can infer state transition rates (arrows). **(B)** Schema for statistical inference of transition rates from clonal inC-RNA-Seq data by fitting stochastic models of cell state-transitions to observed NMF program usage over time. The model was used to infer rates of program induction (*r*_01_) and loss (*r*_10_) for each program in each treatment condition (see Supplemental Text 1). **(C)** Fitted transition rates exemplified for one program (program 3), with all drugs increasing persistence of the ‘high’ state while destabilizing the ‘low’ state. **(D)** Changes in K562 gene expression program persistence times from the fitted dynamic transition rates, showing reduced clonal memory for some programs (blue) but not others (red). **(E)** Corresponding changes in steady-state bias (i.e. the fraction of cells in active state), showing distinct changes from program persistence. **(F)** Changes in persistence are distinct from changes in mean expression for single genes, shown for one drug (Aza) in K562 cells.

As shown schematically in **Fig. 4C**, cells were pre-incubated with Dec, Aza, Vor or vehicle alone at sub-lethal doses for two days (IC30 dosage, see **Fig. S5A** for cell survival curves), then encapsulated and grown in CAGEs over a period of 6 days with continued drug treatment (**Fig. 4C**). At progressive time points, the clones were sampled, lysed in CAGEs and then frozen prior to inC-RNA-seq. In total, we analyzed transcriptomes from 134,805 CAGEs representing single cells sampled at day 0 (n=37,902) and expanded clones grown 2, 4, or 6 days (n=96,903) in different treatment conditions. An equivalent analysis of this number of isolated colonies in wells would require 1,010 96-well plates. In both K562 and L1210 clones, we identified genes with above-Poisson coefficients of variation in gene expression (CV, or standard deviation/mean; shown for untreated cells at days 4 and 6 in **Figs. S5B,C**). The most variable genes between clones defined modules of gene expression that persisted over time, as seen from clustering gene-gene correlations and comparing the results to those from mock clones generated by randomly combining single cell transcriptomes (**Figs. 4D, S5D**). Persistent clonal heterogeneity can also be appreciated by UMAP embedding of the clones (**Figs. S5E**).

To identify persistent gene expression programs varying between clones, we factorized their expression by non-negative matrix factorization (NMF). Approximately 12-15 programs explained variation in gene expression above random for both cell lines (eigenvalue crossover method (*5*), **Fig. S5F**). Several of these were readily identifiable from their gene loadings, including cell cycle-associated programs (K562 programs 12,13 and L1210 program 14; **Figs. 4E, S5G, Table S2**), erythroid differentiation (K562 program 3), putative epithelial (*KRT8/18*-enriched) and mesenchymal (*VIM*-enriched) programs (K562 programs 4, 14) as well as myeloid programs expressing the transcription factors *Irf8* and *Myc,* interleukin receptor *Il7r* (L1210 program 0); and myeloid transcription factor *Klf6*, M-CSF (*Csf1*) and osteopontin (*Spp1*) (L1210 program 7). Significantly, these programs remained variable after clonal expansion beyond that expected from mock clones (**Figs. 4F, S5H**). Several programs also showed mutual exclusive patterns of expression: for example, K562 day 6 colonies were either enriched in program 14 (*VIM*-hi) or program 3 (erythroid, *HBE1/HBA2/ALAS2*-hi) but not both (p<10^-50^, Fisher exact test, adjusted to correct for false discovery), while mock colonies showed mixing of the two programs (**Figs. 4G, S5I,J**).

Next, we explored how Aza, Dec or Vor treatments alter gene expression dynamics. Drug treatment may alter epigenetic inheritance to either bias the fraction of cells that express a program, or the rate at which a program switches states (**Fig. 5A**). To quantify these two effects, we introduced a statistical inference framework that fits the observed distribution of NMF program usages across clones over time to a stochastic model of state switching, parameterized by the persistence times of cells in a ‘low’ state (1/*r_01_*) and ‘high’ state (1/*r_10_*) for each program (**Fig. 5B,C** and Supplementary Text 1). From these fits we obtained changes in program persistence times [the relaxation time-scale to steady-state, 1/(*r_01_*+*r_10_*)] (**Figs. 5D, S6A-C**), and steady-state bias [the fraction of time that a program is active, *r_01_*/(*r_01_*+*r_10_*)] (**Fig. 5E, S6D**). These analyses revealed that all three drugs triggered stereotyped changes in program persistence and bias in K562 cells, with the dynamics of some programs slowing down (higher persistence) and others accelerating. In K562 cells, all three drugs triggered differentiation towards the erythroid state (program 3 high state), with increased persistence in the differentiated state (**Figs. 5C-E**). The changes in persistence times were correlated but distinct from changes in mean expression of individual genes (**Fig. 5F**). In L1210 cells, the DNMT inhibitors (Aza, Dec) showed distinct responses from the HDAC inhibitor (**Fig. S6D**), but again the drugs led to both increases and decreases in state persistence times. The result is that the DNMT and HDAC inhibitors do not reduce clonal heterogeneity over several cell divisions. These observations are consistent with changes in gene expression heritability seen after genetic ablation of DNA methyltransferases(*35*), which previously was done by analyzing individual clones in wells. The specific nature of the effects of these drugs, with a failure to globally reduce persistence, may explain why they sensitize chemotherapy in some patients but not others(*36*). More generally, the approaches taken here for live-cell genomic analysis following perturbations and growth may be generalizable to other colony assays, including on gastruloids and tissue-derived organoids.

## Discussion

In summary, we have established a versatile platform for high-throughput single cell- and single colony-genomic analysis through the development and use of capsules with amphiphilic gel envelopes—CAGEs. The small compartments are biocompatible, quick to produce and have a tunable chemistry. Alternative compositions can be evaluated through biophysical assays to define new permeability, mechanical, adhesive or other functional properties. The approaches reported here for barcoding DNA libraries in capsules (inC-Seq) and to study living cells can be adapted to a range of problems and may be further combined with sorting, fixing and staining steps, each demonstrated here.

As genomic assays become more complex, they face increasing limitations when carried out in droplet emulsions and in fixed cells. CAGEs offer solutions to these limitations, enabling multi-step processing that offers flexibility in developing genomic assays and, importantly, extends functional genomic methods to live-colony and clonal growth assays at scale. Here, we apply CAGEs to scale up an assay of gene expression persistence, which has so far required analysis of individual colonies grown in microtiter plates. We use data from over 100,000 capsules - a scale that allows statistical inference of dynamic transition rates of gene expression programs in cells, and determining rate changes upon treatment with DNMT or HDAC inhibitors. Capsules will likely enable profiling difficult-to-process samples, and they may prove useful in studying perturbation outcomes that depend on cell–cell interactions, such as developmental or immune-cell interactions, and for performing colony-forming assays or studies of organoids at scale. Capsules may also allow evaluation of libraries of signaling molecules, or may be used as vehicles for clonal evolution or selection followed by genomic profiling. These in turn may provide functional information that could support developing predictive, data-driven models of cellular dynamics and interactions.

## Acknowledgments

We thank Linas Mazutis, Denis Baronas and Tim Mitchison for ongoing discussions and feedback during this project, John Oakey for guidance of ROS generation during photopolymerization, the labs of Olivier Pourquié and Fernando Camargo for providing iPSC and HSC cells; Sean McGeary for providing K562 and L929 cells, Yuyang Chen for providing L1210 cells, the lab of Michael Greenberg for access to their Illumina Sequencer; Jeffery A. Nelson at the Bauer Core Facility at Harvard University for FACS work coordination; and the Single Cell Core at HMS for supporting work by TT and AK. The cryo-SEM work was performed in part at the Harvard University Center for Nanoscale Systems (CNS); a member of the National Nanotechnology Coordinated Infrastructure Network (NNCI), which is supported by the National Science Foundation under NSF award no. ECCS-2025158. We extend our heartfelt gratitude to Adam Graham at the Harvard CNS core, whose expertise was instrumental in bringing this cryo-SEM work to fruition. We thank all members of the Klein lab for critical reading and feedback on the manuscript.

## Author contributions

IM developed the capsules and carried out all experiments except FACS and Cryo-SEM. HS supported diffusion assays and optimizing cell culture conditions in capsules. TT carried out flow-sorting in capsules. AK carried out SEM imaging with support of the CNS core facility. IM and AMK conceived and designed experiments and carried out data analysis. AMK supervised the work. IM and AMK wrote the manuscript. All authors provided feedback on the manuscript.

## Data availability

Sequencing data available at Gene Expression Omnibus (GSE306708) and Sequence Read Archive (PRJNA1311090). Python code for Capsule segmentation and pipeline for scRNA-seq read pre-processing will be deposited at github.com/AllonKleinLab/paper-data/tree/master/Mazelis2025_CAGEs.

## Supplementary Materials

**Fig. S1.**
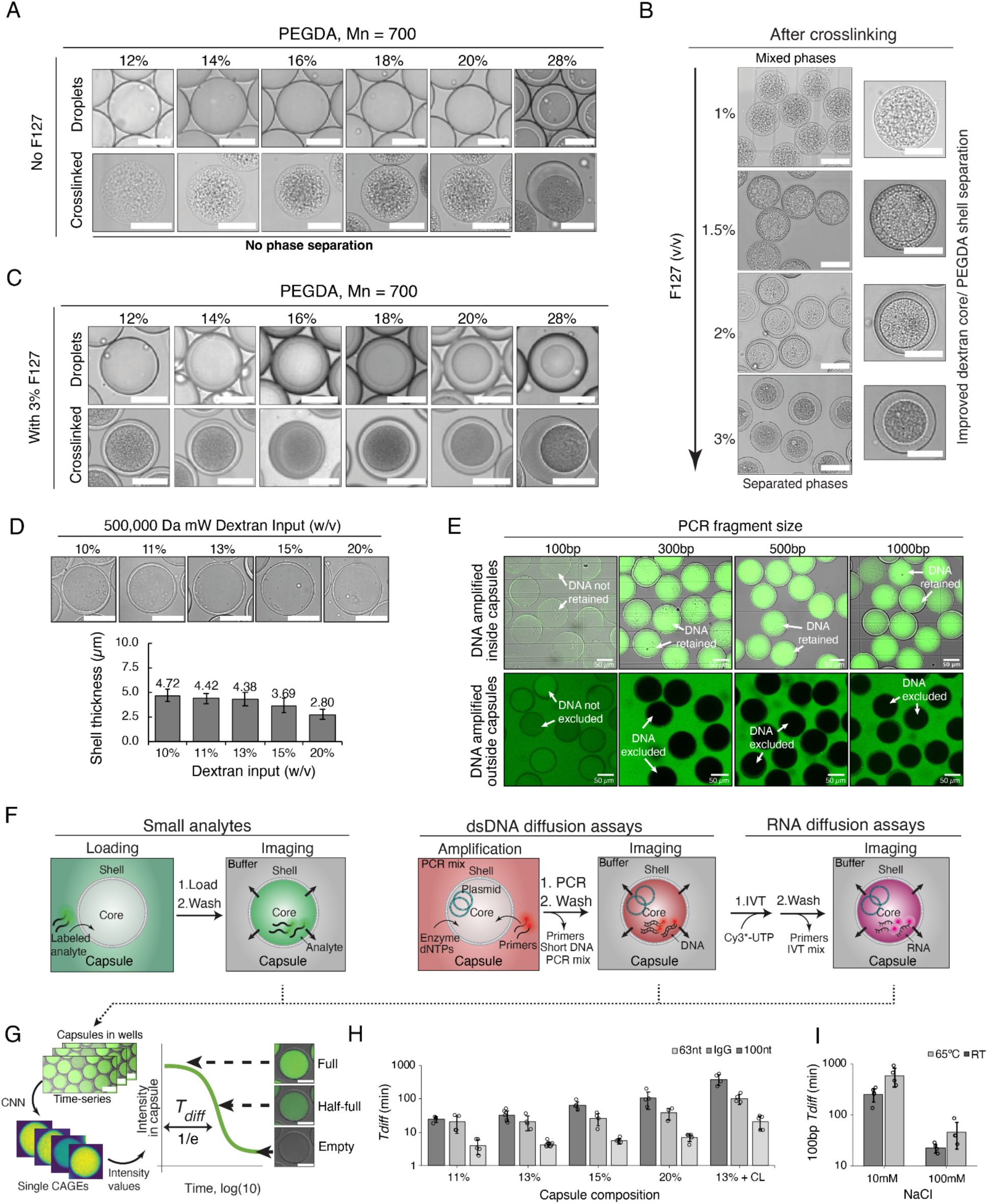
Development and characterization of pluronic F127 diacrylate CAGEs. **(A)** Micrographs demonstrating common failures of hydrogel capsule synthesis using PEG diacrylate as a shell (PEGDA). Capsules fail to form due to the lack of phase separation and form mixed dextran:PEGDA beads, hindering the ability to vary the composition and thus properties of the capsules. **(B)** Micrographs showing successful capsule formation upon addition of Pluronic F127, stabilizing PEGDA:Dextran phase separation with an outer PEGDA phase. **(C)** Same as A, but with addition of 3% F127 to stabilize phase separation, resulting in robust capsule formation across all concentrations evaluated. **(D)** Micrographs showing F127DA capsules (CAGEs) formed using different concentrations of 500 kDA Dextran solution, and quantifying shell thickness. **(E)** Fluorescent confocal microscopy images showing DNA fragments ≥ 300bp selectively retained (top) or excluded (bottom) by F127DA CAGEs. DNA fragments of different sizes were amplified for 16 cycles from a plasmid DNA template encapsulated inside the capsules (top) or added to the outside of empty CAGEs (bottom). Top row samples were washed to remove background signal. Samples were stained using SYBR Safe DNA dye. **(F)** Experimental schematics for measuring transport rates of different analytes through the capsules shell. Rapidly moving, small analytes (66nt, 100nt, IgG) were loaded into the capsules from the outside buffer, followed by washing and imaging (left). dsDNA that’s too large to quickly enter capsules was loaded using PCR by amplifying 100, 300, 500, 1000bp DNA with labeled primers of an encapsulated plasmid DNA, followed by washing and imaging (middle). 300/500/1000nt RNA molecules were generated in CAGEs by carrying out an IVT reaction, using amplified DNA harboring T7 promoter as template (right). **(G)** Schematic of time-lapse data collection to measure diffusion rates. Images were processed by segmenting capsules from each field of view, obtaining average core fluorescence intensities over time, fitting with double exponential decay and extracting time-scales as time required to reach 1/e of the initial intensity. **(H)** Diffusion half-times are tunable by generating CAGEs with different dextran concentrations, and with optional addition of 2% PEG4DA crosslinker (CL). Higher values = slower diffusion through the capsule shell. See legend for the three analytes evaluated. Plot represents mean data from ≥3 independent experiments with SD as error bars. **(I)** Diffusion half-times in different concentrations of salt and different temperatures. Increased buffer temperature and lower salt concentration both slow diffusion. All scale bars = 50 µm. Plot represents mean data from ≥3 independent experiments with SD as error bars.

**Fig. S2.**
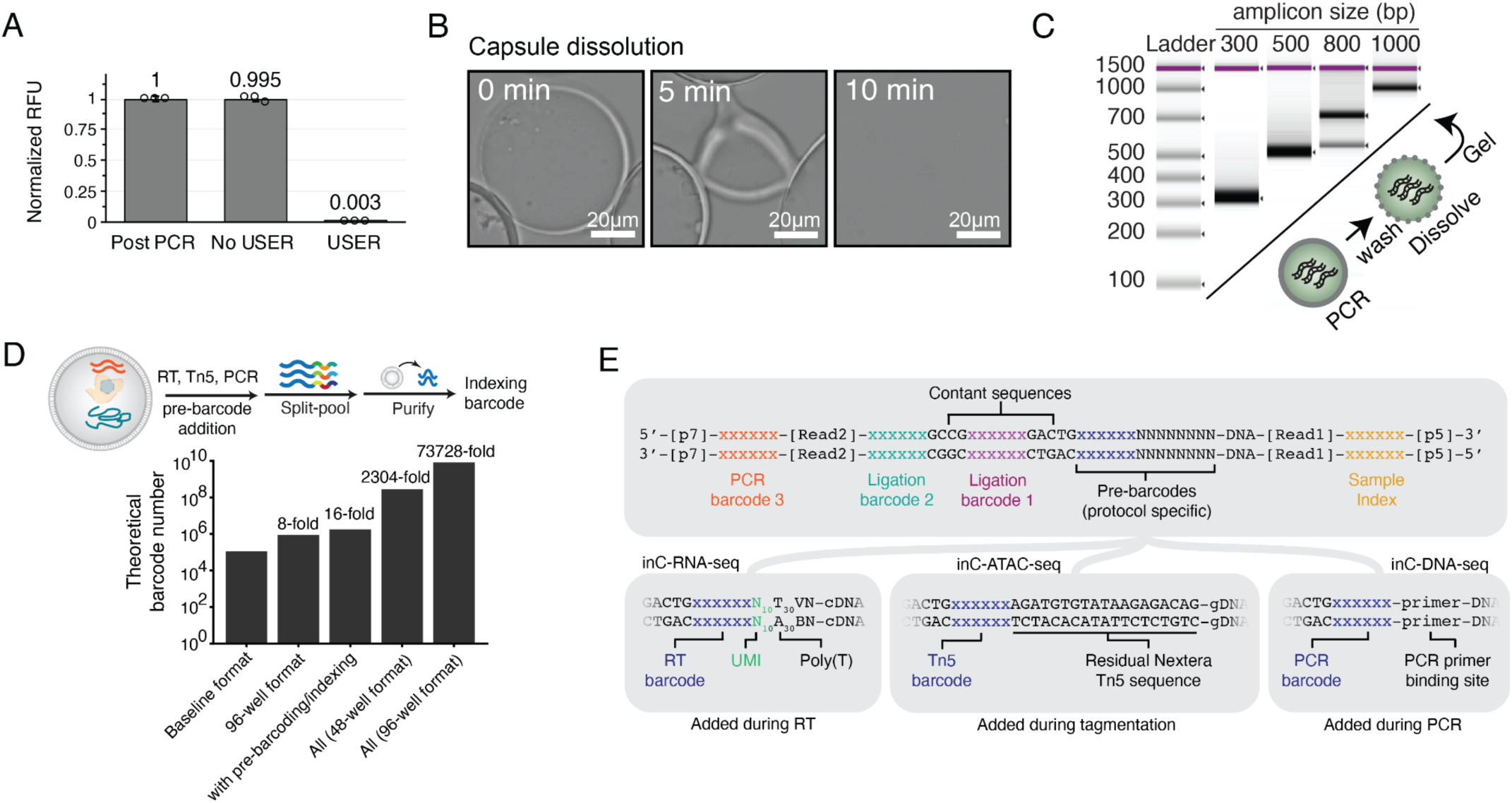
Characterization of capsule library processing, scalability and library structure of inC-Seq. **(A)** Quantification of dU excision in CAGEs using USER enzyme. DNA, amplified in capsules using Cy5 labeled dU harboring primer is retained inside the CAGEs as seen from fluorescent imaging (Post PCR). Addition of USER enzyme efficiently excises dU (and thus Cy5) as quantified by Cy5 signal loss (USER). Plot represents mean data from 3 independent experiments with SD as error bars. **(B)** Micrographs showing CAGEs dissolved in the presence of 1M NaOH **(C)** DNA purified from dissolved CAGEs, analyzed using a BioAnalyzer. Each lane corresponds to capsules with PCR amplicons of a defined size. **(D)** Theoretical barcode space of inC-seq, giving realistic designs from ∼110K to ∼8B barcodes. A baseline is to carry out 3 rounds of barcoding, with 48 barcodes per round (number barcodes = 48^3^). This can be scaled by increasing the plate size format or including additional rounds of barcode addition with a specific number of barcodes during RT, ATAC or pre-PCR steps (pre-barcoding) and after dissolving capsules in batches (indexing). **(E)** The library structures for inC-seq, showing protocol specific barcodes incorporated during RT (barcode and UMI), tagmentation and PCR. Usage of dU, enables barcode addition between Tn5 cut site and Illumina read primer binding site.

**Fig S3.**
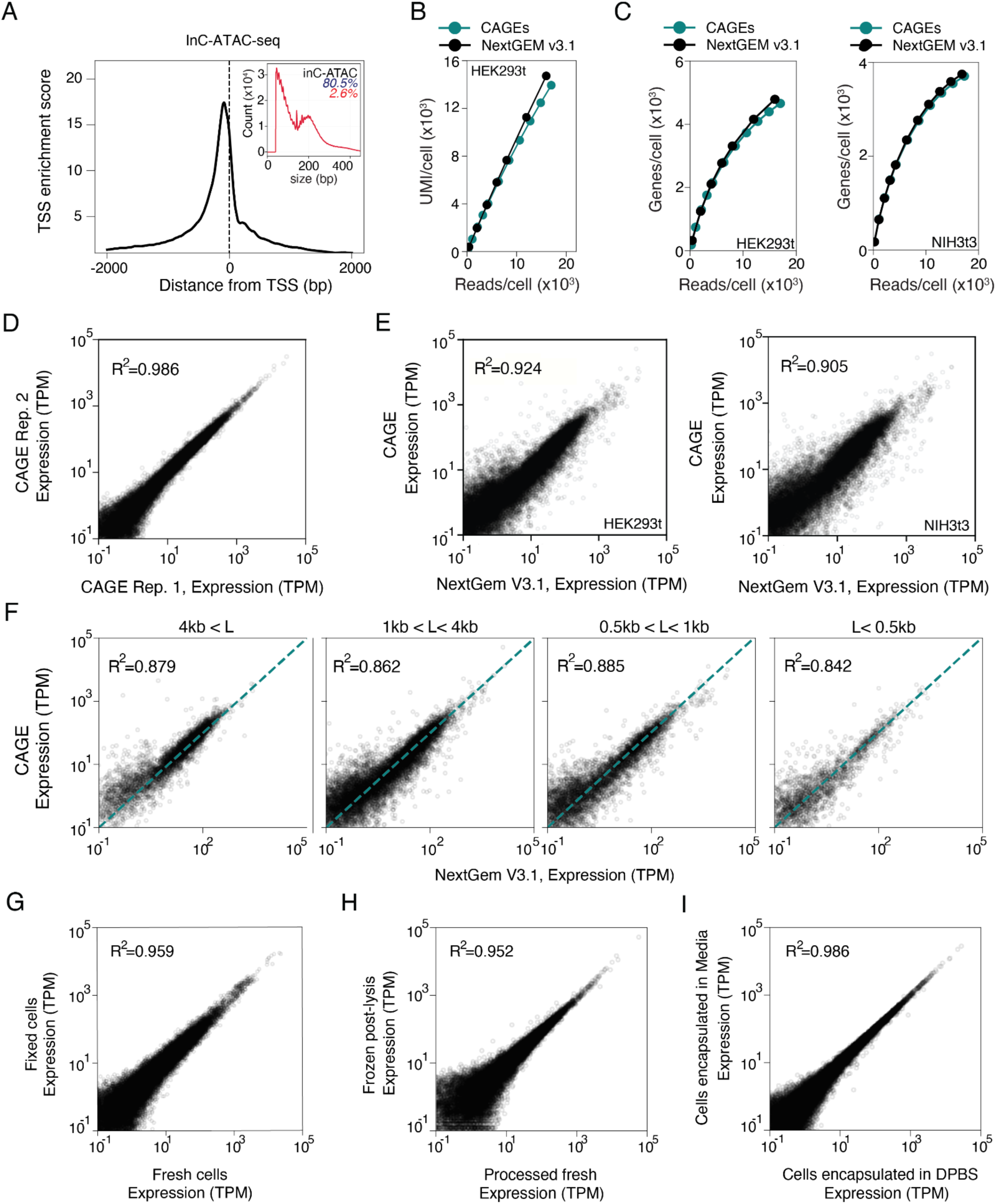
Additional technical benchmarking for inC-ATAC-Seq and inC-RNA-Seq. **(A)** Capsule inC-ATAC-seq data shows high Transcription Start Site (TSS) enrichment. Inset shows DNA library fragment size distribution. Blue text - % reads mapping to combined mouse and human reference genomes, red text - % reads mapping to mitochondrial DNA. **(B)** UMI/read plots following downsampling reads from scRNA-Seq libraries generated in capsules (inC-RNA-seq) or using publicly available 10× Genomics NextGem v3.1 data, showing comparable library complexity in HEK293t samples. **(C)** Gene/read detection plots following downsampling reads from scRNA-Seq libraries generated in capsules (inC-RNA-seq) or using publicly available 10× Genomics NextGem v3.1 data, showing the same detection level in HEK293t and NIH3t3 samples. **(D-E)** Gene expression comparisons, showing high technical reproducibility of inC-RNA-seq **(D)** and agreement with publicly available 10× Genomics NextGem v3.1 data in HEK293t and NIH3t3 cells **(E)**. **(F)** Capsules do not show bias toward captured transcript length as depicted by gene-expression agreement with 10× Genomics NextGem v3.1 at different gene length cutoffs. **(G, H, I)** Comparison of mean gene expression of inC-Seq libraries between fresh vs. PFA-fixed cells **(G)**; immediate analysis vs in-capsules cell lysate cryopreservation **(H);** and following cell encapsulation in tissue culture media vs PBS **(I)**.

**Fig S4.**
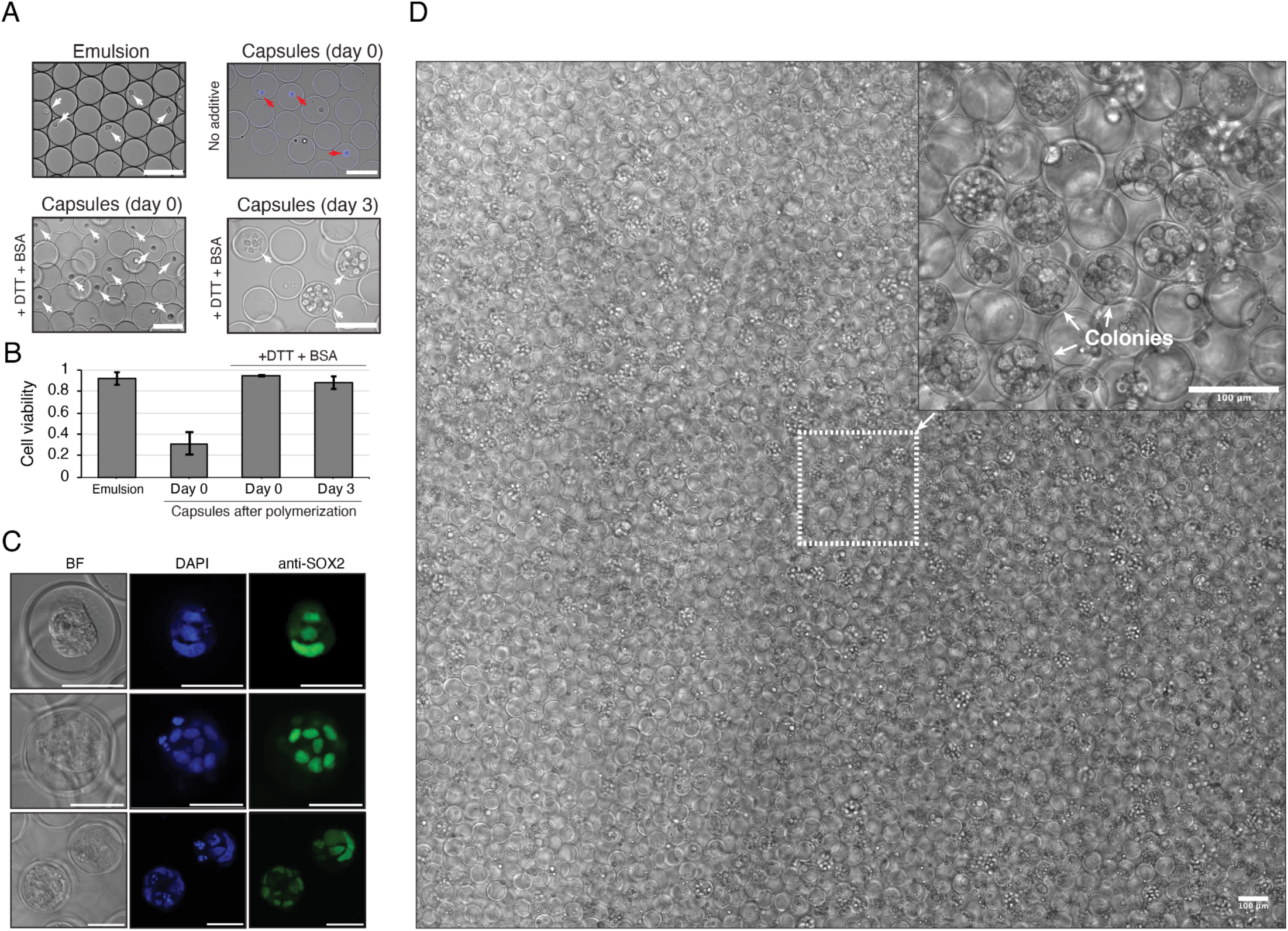
Live cell culturing and colony expansion in CAGEs. (A, **B)** Optimization of encapsulation conditions for live cell culturing. **(A)** Cells, encapsulated into droplets (top left), readily die after CAGE polymerization (top right) due to ROS generation. Addition of 8 mM DTT and BSA during CAGE generation protects the cells from ROS resulting in high viability (bottom left) and enables colony formation (bottom right). (B) Quantification of cell viabilities. Plot represents mean data from 3 independent experiments with SD as error bars. **(C)** Micrographs showing clones derived from single human induced pluripotent stem cells (hiPSCs) following 4 days in culture and in-capsule fixation and immunostaining for SOX2. Scale bar = 50µm **(D)** Micrograph of high-density K562 colonies expanded from 1-4 cells in CAGEs cultured in a culturing 12-well plate. Scale bar = 100µm.

**Fig S5.**
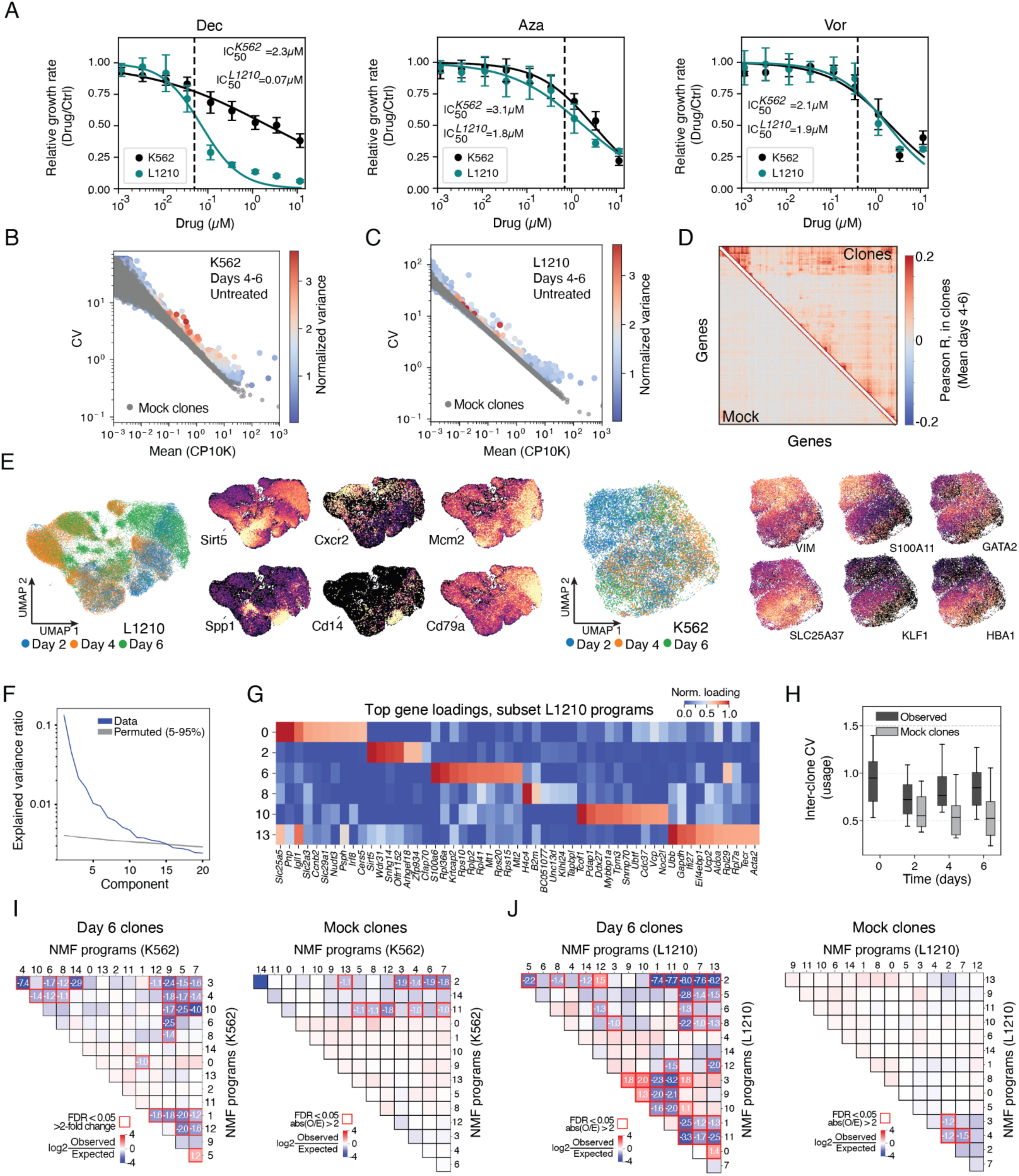
Analysis of gene expression persistence in clones by inC-Seq. **(A)** Dose response curves for K562 and L1210 cells for the three drugs used to select dosages for experiments in **Figs. 4-5**, showing growth rate normalized to control. Growth rate = log[(cell number at day 4)/(cell number at day 0)]/(4 days). Data shows mean values with SEM of n=6 (Dec, Aza) and n=4 (Vor) replicate experiments. IC50 values are indicated, and dashed lines indicate the drug concentrations selected for subsequent analysis (approximately IC30). **(B,C)** Gene coefficients of variation (CV)-mean plots for the two cell lines in control clones, averaged across 3 replicate samples and two time points (days 4,6), and overlaid with values from mock clones generated by randomly combining single cell transcriptomes from day 0 (see methods). CV values were calculated separately within each (sample, timepoint) pair and then averaged across samples, weighted by the number of clones per sample. Colorbar gives the corresponding weighted-average normalized variance (see methods). **(D)** Gene-gene correlation for L1210 cells, corresponding to **Figs. 4D**. **(E)** UMAP representation of gene expression heterogeneity between L1210 and K562 colonies. **(F)** Variance explained by top principal components in K562 cells, compared to randomized data with the same marginal gene expression distribution per gene. The cross-over occurs at 15 components. **(G,H)** L1210 plots corresponding to **Figs. 4E,F**. **(I,J)** Heatmaps showing multiple gene expression programs are mutually exclusive in (I) K562 and (J) L1210 day 6 clones, as seen from low numbers of clones observed mixed NMF programs as compared to expectation (Observed/Expected = *f_XY_/f_X_f_Y_*). Left panels: real data; right panels: plots generated from mock clone data, showing loss of persistent clonal variation.

**Fig S6.**
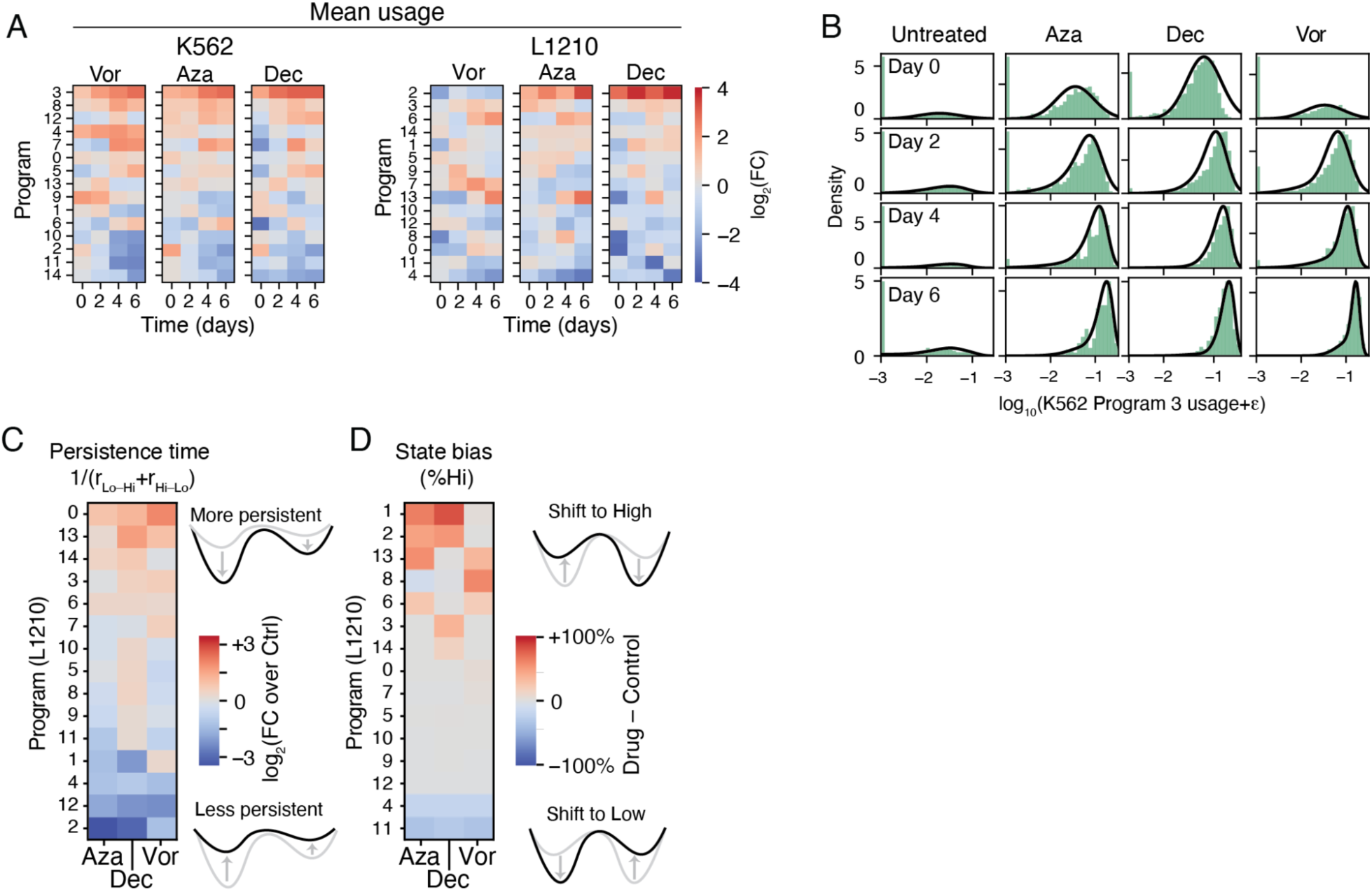
DNMT and HDAC inhibitors alter persistence and bias in gene expression program activation states for cells grown in CAGEs. **(A)** NMF Program mean usage dynamics in K562 and L1210 cells after drug treatment relative to controls. **(B)** An example of the fits used to infer state switching rates from the model defined in Fig. 5B and Supplementary Text 1. Green bars show histograms of observed NMF usage, here for K562 NMF program 3 (erythroid program). Black curves show the model fits. **(C,D)** Changes in inferred L1210 NMF Program persistence time and bias relative to controls. Corresponding to Fig. 5D**,E**.

## Materials and Methods

### CAGE synthesis

Capsules with amphiphilic gel envelopes (CAGEs) were prepared using a co-flow droplet microfluidic PDMS device (Darwin Microfluidics, DG-MCN-C4) by flowing shell and core solution at 170µL/h into fluorinated HFE7500 carrier phase supplemented with 2.5% (w/v) 008-FluoroSurfactant (Ran biotechnologies) infused at 580µL/h. Shell polymer solution consisted of 8% (v/w) F127DA (Creative PEGWorks, PPO-121-5g), 1% (w/v) PPPDA (Sigma Aldrich, 929611-500MG) in 1x DPBS (Thermo Fisher Scientific, 14190136), unless stated otherwise. The core solution consisted of a 13% (w/v) dextran (mW = 500,000Da) (Sigma Aldrich), 0.1% (v/v) of F68 (Thermo Fisher, 04196546SB) and 0.2% (w/v) of LAP (lithium phenyl-2,4,6-Trimethylbenzoyl phosphinate) (Sigma Aldrich, 900889-1G)] in 1x DPBS, unless stated otherwise. Generated droplets were collected on ice into 1.5mL Eppendorf tube unless stated otherwise. After collection, the tube was incubated on ice for 5 min, followed by incubation at 30°C for 3 mins and removal of bottom HFE7500, unless stated otherwise. Next, the tube was hand shaken 5 times and incubated at 30°C for additional 2 mins. The tube was carefully transferred to a bottomless tube holder and exposed to 405 nm light from below for 35s, unless stated otherwise. After polymerization, hydrogel capsules were purified by removing the bottom HFE7500 phase, adding 100µL 20% (v/v) PFO (Sigma Aldrich, 370533-25G) in HFE7500, shaking, and spinning down for 5s. One additional wash with HFE7500 was performed after removal of PFO/HFE7500 followed by three 5s spins to remove any remaining HFE7500 droplets. The resulting hydrogel capsule pellet was resuspended in a wash buffer as described below. PEG-shell capsules were generated as CAGEs, but using PEGDA instead.

### CAGE washing

Capsules were washed by resuspending 100-300µL of packed capsules in 800-1000µL of wash buffer, vortexing/pipetting and spinning down at 300g - 800g for 30-60s (depending on the application). The supernatant is then aspirated leaving ∼20% of volume behind to not disturb the packed capsules. Different capsule wash buffers were used as described below with corresponding composition: TI (10 mM Tris, 0.1% Igepal CA-630 (Millipore Sigma, I8896-50ML)); TI10 (10 mM Tris, 10mM NaCl, 0.1% Igepal CA-630); TI100 (10 mM Tris, 100mM NaCl, 0.1% Igepal CA-630); TIbS (10 mM Tris, 0.1% Igepal CA-630, 0.05% SDS); TEIS (10 mM Tris, 0.1mM EDTA, 0.1% Igepal CA-630, 0.1% SDS); HI10 (Nuclease free water, 10mM NaCl, 0.1% Igepal CA-630), 3xSSCI (3x Sodium saline citrate buffer (pH 5.9) (Thermo Scientific), 0.1% Igepal CA-630).

### Cryo-SEM imaging of CAGEs

CAGEs were stored in TI containing 10mM Tris pH8, 0.1% Igepal prior to imaging. A 10μl sample was loaded onto each of three gold planchets (Leica #16770132; Carrier, Dia. 3.0×0.8mm, Cu-Au, dome) and rapidly frozen using slushy nitrogen (made using Edwards Plate Degasser PD3; Model RV3, Serial No. 056188208, Code No. A652-01-903) and transferred to the Leica EM VCM loading station (SN 614252, #16771611104). The positioned frozen planchets were next transferred from the Leica EM VCM loading station to the Leica ACE 600 sample preparation chamber (SN 601238, #16771525) with the help of the Leica EM VCT 500 vacuum cryo-transfer system maintained at - 150°C using liquid nitrogen. In the chamber, the samples were knife-fractured at -150°C, sublimated at -105°C for 9 minutes and 10 seconds, then re-cooled to -150°C. The samples were sputter-coated with a 10nm layer of Platinum/Palladium (80:20) and transferred to the Zeiss Gemini 360 <u>F</u>ield <u>E</u>mission <u>S</u>canning <u>E</u>lectron <u>M</u>icroscope (FESEM) (GeminiSEM360 349599-9100-010, SN 8217010169), with the help of the VCT 500 vacuum cryo-transfer system. Imaging was performed at -150°C on a temperature-controlled FESEM stage at 3 keV using the Everhart Thornley detector.

### Analyte cross-shell diffusion assays

Rates of analyte diffusion through the capsule shell were assessed using time-lapse confocal microscopy (Fig. S1G). Fluorescent analytes, loaded into capsules as described below, generate a bright signal in the capsules core that decreases over-time as molecules diffuse out into the non fluorescent surrounding buffer. Capsules housing different analytes in different buffers were imaged at 20× magnification in 96-well glass bottom plates (MatTek, P96G-1.5-5-F) until the fluorescence signal from the core decreased below 1/e of initial value but no longer than 14 days. Acquired time-lapse movies were processed by segmenting capsules from each field of view using a custom PyTorch convolutional neural network (github.com/AllonKleinLab/) and average core fluorescence intensities were extracted. Average fluorescence time-series were fitted with double exponential decay and time-scales were extracted as time required to reach 1/e of the initial intensity. Values for analyzed buffers and analytes are provided in **Table S1**.

### Small analyte assay

Cross-shell diffusion half-lives of 63-100 nt ssDNA and IgG were measured by first incubating empty capsules in a buffer containing the analytes (Fig. S1F left). For this step, empty hydrogel capsules were washed 3 times with TI100 (pH 8.0) buffer and ∼5µL of packed capsules were transferred to 1.5mL tubes. 1µL of fluorescently labeled analyte solution (10µM Cy5-63nt; 10µM FAM-100nt [**Table S3**, diffusion], 5 mg/mL FITC Anti-mouse CD45 IgG (BioLegend, 110705) was added on to 5µL of packed capsules and left on ice for 2 hours. After incubation, capsules loaded with analytes were washed with TI (pH 8.0) buffer twice followed by a single wash with a buffer containing different salt concentrations (TI, TI10, TI100). Capsules were packed, transferred to a 96 well glass bottom plate and resuspended to a final volume of 40µL in a buffer of interest.

### dsDNA diffusion assay

dsDNA that is too large to quickly load into capsules by diffusion was generated within-capsules using a PCR reaction (Fig. S1F middle). First, capsules were generated following the normal procedure, with a plasmid DNA added into the dextran core mix at a final concentration of 10pg/mL prior to encapsulation. Capsules were polymerized and washed three times with TI100 (pH 8.0), twice with TI10 (pH 8.0) and twice with TI (pH 8.0). 12.5µL of packed capsules were used in a 25µL PCR reaction consisting of 2.5µL of 10µM universal Cy3 labeled primer, 2.5µL of 10 µM of Xbp_RV primer [**Table S3**, diffusion], 1µL of 10mM of dNTPs (NEB, N0447L), 5µL of 5x Phusion HF reaction buffer (NEB, E0553L), 1µL of 10% Igepcal CA-630 and 0.5µL of Phusion polymerase (NEB, E0553L). Here, X=100, 300, 500 and 1000 corresponds to primers generating different size amplicons (see **Table S3**, diffusion). DNA was amplified for 15 cycles using the different primers, to generate dsDNA amplicons of length 100/300/500/1000bp. The reaction was carried out by first incubating the tubes on ice for 5 mins, followed by a PCR cycling program: 98°C for 30s; 15 cycles of 98°C for 10s, 65°C for 20s, 72°C for 10s (for 100bp and 300bp products) or 30s (for 300bp and 1000bp products); and a final extension of 72°C for 5 min and hold at 25°C. After PCR, capsules were washed twice with TI100 (pH 8.0), twice with TIbS (pH 8.0), twice with TI10 (pH 8.0), twice with TI (pH 8.0) and once with buffer of interest. Capsules were packed, transferred to a 96 well glass bottom plate and resuspended to a final volume of 40µL in a buffer of interest.

### ssRNA diffusion assay

300/500/1000nt RNA molecules were generated in capsules by carrying out an IVT reaction (Fig. S1F right). First, dsDNA of appropriate size was amplified in capsules of a plasmid DNA template as described above by carrying the PCR reaction for 8 cycles with a primer housing a T7 promoter [**Table S3**, diffusion]. After PCR, capsules were washed twice with TI100 (pH 8.0), twice with TIbS (pH 8.0), twice with TI10 (pH 8.0), twice with TI (pH 8.0). IVT was carried out in 21µL reactions, consisting of 1X T7 reaction buffer (NEB, M0251S), 8mM of ATP, GTP, CTP (NEB, N0466L), 6 mM of UTP (NEB, N0466L), 0.16mM of Cy3-UTP (APExBIO, B8330), 4U of T7 polymerase (NEB, M0251S) and 1.06U RiboLock RNAse Inhibitor (ThermoFisher, EO0381) for 60 min. After IVT, capsules were washed twice with TIbS (pH 8.0), twice with TI100 (pH 8.0), twice with TI (pH 8.0) and once with the buffer of interest. Capsules were packed, transferred to a 96 well glass bottom plate and resuspended to a final volume of 40µL in a buffer of interest.

### CAGE flow cytometry

FITC fluorescent capsules used for optimization of fluorescence-activated capsule sorting (FACS) were synthesized as described above, except Fluorescein isothiocyanate-dextran (FITC-dextran) (ThermoFisher) was added in to the core mix at 169 µg/mL. Non-fluorescent capsules were synthesized as above. After capsule polymerization and HFE7500 removal, capsules were washed once with 300µL of hexane. The capsules were counted by pipetting 10 µL of a 1:10 dilution of stock capsules into a 96-well plate to determine concentration of capsules prior sorting sample preparation. FITC+ and FITC-capsules were mixed so that the final fraction of FITC+ was 0.05-0.1. Capsules were transferred to a 15 mL conical tube and spun down at 800 g for 1 minute. Supernatant was removed and capsules were resuspended in DPBS + 0.1% Igepal or TI to a final concentration of 500 – 1,000 capsules/µL in the sample tube via pipetting. Capsules were sorted using a 200 µm nozzle on the Beckman Coulter MoFlo Astrios EQ high speed cell sorter. “Enrich” or “Purify” sort modes were used in addition to a “1-2 Drop” Drop Envelope. Drop Delay values were manually calibrated using FlowCheck beads (Beckman Coulter, 6605359). The Drop Delay value determined using the FlowCheck beads was offset by +/-0.5 units. To check Drop Delay calibration using capsules, 50 capsules were sorted onto a glass slide and quickly assessed for purity and recovery using fluorescence microscopy. Once calibrated, capsules were sorted into 1.5 mL Eppendorf tubes (Eppendorf). After sorting, the collection tubes were rinsed with DPBS + 0.1% Igepal or TI buffer to dilute the sheath fluid and dislodge capsules from tube walls. Sorted sample purity and recovery were assessed by sampling the capsules and imaging them on a Nikon Ti1000 wide-field fluorescent microscope. Purity and recovery were estimated by counting the fluorescent and non-fluorescent capsules and estimating the number of total capsules using the number of counted capsules and volume of the sorted sample.

### CAGE dU PCR and USER excision

A 50µL volume PCR reaction utilizing a dU-containing primer (dU PCR) was carried out as follows: a reaction mix was assembled consisting of 10µL of 5X Q5U reaction buffer (NEB, M0515L), 2µL of 10mM dNTPs (NEB, N0447L), 5µL of 10µM FW primer (protocol specific), 5µL of 10µM dU universal RV primer [**Table S3**], 25µL of packed capsules, 1µL of Q5U High-Fidelity DNA polymerase (NEB, M0515L), and 2µL of 10% Igepal CA-630 (Millipore Sigma, I8896-50ML) unless stated otherwise. To set up the reaction, capsules were transferred to PCR tubes on ice and the rest of PCR reagents were added ensuring proper mixing by pipetting. The reaction was carried out by first incubating the tubes on ice for 5 mins, followed by a PCR cycling program: 98°C for 30s; 6 cycles of 98°C for 15s, 68°C for 20s, 72°C for 4 min; a final extension of 72°C for 2min and hold at 25°C, unless stated otherwise. After the cDNA amplification, capsules were washed three times with TI100 (pH 8.0), twice with TIbS (pH 8.0), three times with TI100 (pH 8.0) and twice with TI10 (pH 8.0). Capsules were imaged on a widefield microscope. For dU excision, a reaction was performed in TI10 (pH 8.0) buffer with 0.01U/µL of USER^TM^ enzyme (NEB, M5505L) in a shaking heat block at 37°C for 20 min. Following the reaction, capsules were washed twice with TI100 (pH 8.0), once with TIbS (pH 8.0) and TI10 (pH 8.0). To fully remove the cleaved DNA sequence, capsules were washed in a 1 mL of TI (pH 8.0) buffer with a 5 min incubation at 60°C, twice, with a room temperature TI (pH 8.0) wash in between. Capsules were imaged on a widefield microscope.

### CAGE split-and-pool barcoding

Barcoding is carried out through several steps:

#### 1. USER digestion and washes

dsDNA barcoding in capsules was carried after a dU PCR reaction as described above. All protocols described in this paper utilized a 5’CGTGATGCTGTACTTACATGTGACdUG-’3 PCR handle.

Following dU PCR, the capsules were transferred to a 1.5mL tube and dU was excised and washed as described above. The capsules were then washed twice with TI100 (pH 8.0), once with TIbS (pH 8.0) and TI10 (pH 8.0). To fully remove the cleaved DNA sequence, capsules were washed as described above, with one modification: in the final two washes using TI (pH 8.0), the buffer was supplemented with 200mM MgCl2.

#### 2. Preparation of barcoding plates

Work done in this paper used a 48-barcode format. Plates containing Ligation-1 and Ligation-2 barcodes were prepared in advance by mixing 100µM forward and reverse oligos (**Table S3**, Ligation-1, Ligation-2) at 1:1 ratio. Barcodes were annealed by heating the plates at 95°C in a PCR machine for 5 min and then letting them plate cool at room temperature for 1 hour, followed by incubation on ice. Annealed barcoding plates were kept frozen at -20°C. A third plate for PCR-3 barcoding was also prepared in advance, with wells each containing 10µmM barcoding primers (**Table S3**, PCR-3). Plates were kept frozen at -20°C.

#### 3. Ligation-1 barcode addition

Capsule barcode-1 ligation was performed in a 96-well plate. Reaction volumes were scaled based on the volume of capsules being barcoded as follows. For every 2µL of capsules/well, a reaction is carried out in 10 µL volume with: 5 µL of 2x StickTogether™ DNA Ligase Buffer (NEB, M0318L), 2 µL of capsules in TI with 200mM MgCl2, 2 µL of 50µM annealed barcode duplex and 1µL of T7 DNA ligase. (NEB, M0318L). Before setting up the reaction, capsules were pelleted in TI (pH 8.0) with 200mM MgCl2 and ∼80% of the buffer was aspirated. The amount of buffer to leave behind is determined by rounding the packed capsule volume to a convenient to work volume (for example, 170µL of packed capsules would be aspirated to 200µL using another 1.5mL tube loaded with buffer as visual standard). Packed capsules were left on ice for at least 5 min. A fresh Multiplate^TM^ 96-well PCR plate (BioRad, M0318L) was placed on ice and 1/50th of total capsule volume was transferred to each well using a pipette (in the case of 200µL, 4µL would be added to each well). Next, a ligation master mix composed out of 2x StickTogether™ DNA Ligase Buffer (NEB, M0318L) and T7 DNA ligase (NEB, M0318L) was prepared for 50 reactions in the same tube that housed the aliquoted capsules and 1/50th of the volume was added directly to capsules in each well without pipetting. Finally, thawed and vortexed ligation-1 DNA barcodes were added to wells housing capsules and ligation mastermix using a 12-tip multichannel pipette (volume defined above), followed by pipetting with the same tip to mix the reagents within each well. The barcoding plate was spun down, covered with a silicone mat (Millipore Sigma, Z374938-10EA) and incubated on a heat block at 25°C for 30 min.

After incubation, the reaction was stopped and capsules were pooled by adding 100µL of TIbS (pH 8.0) into each well and then transferring the volume from all wells into a 15mL falcon tube. Each well was washed twice and the plate was inspected under the inverted microscope to make sure no capsules are left behind. Capsules were transferred to a 1.5mL tube and were washed four times with TI10 (pH 8.0).

#### 4. Overhang generation by dU excision

Next, DNA overhangs for Ligation-2 barcode addition were generated by performing a dU excision reaction for 20 min in TI10 (pH 8.0) at 37°C in the presence of 0.01U/µL of USER (NEB, M5505L). After the reaction, capsules were washed twice with TI10 (pH 8.0) and twice using TI (pH 8.0) with 200mM MgCl2.

#### 5. Ligation-2 barcode addition

Capsule Ligation-2 barcode addition was performed in a 96-well plate by repeating the steps described in *Step 3* above.

#### 6. PCR-3 barcode addition

Capsule PCR-3 barcode addition was performed in a 96-well plate. Reaction volume scales with the volume of capsules being barcoded. For 1-2.6µL of capsules, a standard 10 µL reaction houses 5 µL of 2x NEBNext® High-Fidelity 2X PCR Master Mix (NEB, M0543L), 0.4µL of 10% Igepal CA-630 (Millipore Sigma, I8896-50ML), 1-2.6 µL of capsules in TI (pH 8.0), 1µL of 10µM universal PCR primer, targeting the other end of DNA molecule, 1 µL of 10µM PCR barcode (**Table S3**, PCR-3) and water up to 10 µL. Before setting up the reaction, capsules were pelleted, resuspended and distributed over the wells of the barcoding plate as described in *Step 3*. Next, a PCR master mix, composed out of 2x NEBNext® High-Fidelity 2X PCR Master Mix (NEB, M0543L), Igepal CA-630 (Millipore Sigma, I8896-50ML), universal PCR primer and water (if used). was prepared for 50 reactions in the same tube that housed the aliquoted capsules and 1/50th of the volume was added directly to capsules in each well without pipetting. Finally, thawed and vortexed PCR-3 DNA barcodes were added to wells housing capsules and PCR master mix as described in *Step 3,* using a 12-tip multichannel pipette (volume defined above). The reaction plate was spun down, covered with adhesive PCR film (ThermoFisher, AB0558), incubated on ice for 5 min and followed by a PCR cycling program: 98°C for 45s; 5 cycles of 98°C for 10s, 68°C for 20s, 72°C for 4 min; a final extension of 72°C for 2min and hold at 25°C.

After the PCR-3 barcode addition, capsules were pooled by adding 100µL of TIbS (pH 8.0) into each well and transferring all the volume into a 15mL falcon tube. Each well was washed twice and the plate was inspected under the inverted microscope to make sure no capsules are left behind. Capsules were transferred to a 1.5mL tube and were washed three times with TI100 (pH 8.0), three times with TI10 (pH 8.0) and twice with TI (pH 8.0). At this point, capsules were split into 1.5mL tubes for DNA purification and further downstream processing.

### CAGE dissolution and DNA purification

Capsules were dissolved in the presence of 0.5M NaOH, 0.5% Triton X-100 and 0.5% SDS in a shaking heat block set to 40°C for 20min in 1.5mL tubes. The solution was neutralized by adding an equimolar amount of HCl and 200mM Tris (pH 8.0) buffer to a final volume of 100µL. DNA from solubilized capsules was extracted using Qiagen MinElute DNA purification kit (Qiagen, 28004) or AMPure XP magnetic beads (Beckman Coulter, A63881).

### Post-purification barcoded DNA amplification

Barcoded and purified DNA was amplified using PCR. A standard 50µL reaction was composed out of 25µL 2x NEBNext® High-Fidelity 2X PCR Master Mix (NEB, M0543L), 2.5µL of 10µM P7_rv primer (**Table S3**), 2.5µL of 10µM FW primer (protocol depended), 20µL of purified barcoded DNA. Reactions were performed using a PCR cycling program: 98°C for 30s; 10 cycles of 98°C for 15s, 68°C for 20s, 72°C for 2min; a final extension of 72°C for 2min and hold at 4°C unless stated otherwise.

### Library indexing PCR

Indexing PCR was carried out by mixing purified DNA with 25 μL of 2x NEBNext® High-Fidelity 2X PCR Master Mix (NEB, M0543L) and 2.5 μL of 20µM P7_rv, 2.5µL of 20µM indexing primer that houses an 8bp library index (**Table S3**, indexing) and water to 50µL. PCR cycling program: 98 °C, 2 min; 3 cycles of 98 °C, 20 s; 55 °C, 30 s; 72 °C, 20 s and 9 cycles 98 °C, 20 s; 65 °C, 30 s; 72 °C, 20 s; with a 1 min final extension at 72 °C, and 4 °C, hold at the end.

### Plasmid DNA sequencing

Hydrogel capsules containing plasmid DNA were generated as described in *Hydrogel Capsule synthesis*, except the core solution contained 1ng/µL of plasmid DNA. Capsules were polymerized and washed three times with TI100 (pH 8.0), twice with TI10 (pH 8.0) and twice with TI (pH 8.0). Next, 14 reactions of capsule PCR were performed in a 50 µL volume [10µL 5X Q5U reaction buffer (NEB, M0515L), 2µL of 10mM dNTPs (NEB, N0447L), 2.5µL of 20µM GFP/RFP FW primer (**Table S3**), 2.5µL of 20µM dU barcoded CMV [1-14] RV primer (**Table S3**, PCR), 27.5µL of packed capsules, 1µL of Q5U High-Fidelity DNA polymerase (NEB, M0515L), 2µL of 10% Igepal CA-630 (Millipore Sigma, I8896-50ML), 2.5µL of water]. The reaction was carried out by first incubating the tubes on ice for 5 mins, followed by cycling: 98°C for 30s; 14 cycles of 98°C for 15s, 68°C for 20s, 72°C for 4 min; a final extension of 72°C for 2min and hold at 25°C. After PCR, capsules were washed once with TIbS (pH 8.0), three times with TI100 (pH 8.0) and three times with TI10 (pH 8.0). The capsules were then subject to a second PCR reaction performed in 50µL volume as described above, except 2.5µL of 20µM GFP/RFP FW, 2.5µL of labeled dU universal RV primer and 30µL of packed capsules with no extra water were used using the same cycling conditions for 8 cycles. After the DNA amplification, capsules were washed three times with TI100 (pH 8.0), twice with TIbS (pH 8.0), three times with TI100 (pH 8.0) and twice with TI10 (pH 8.0). Washed capsules were subjected to split-and-pool barcoding as described above.

Following split-and-pool barcoding, capsules were split into 16 multiple 1.5mL tubes and were dissolved as described above. DNA was purified using MinElute PCR purification kit (Qiagen, 28004) and barcoded DNA was eluted into 20µL of the elution buffer. Barcoded and purified DNA was amplified using PCR as described in *Post-purification barcoded DNA amplification*, using 2.5µL of TruSeq_R1_GFP/TruSeq_R1_RFP or TruSeq_R1x_GFP/TruSeq_R1x_RFP primer mix (**Table S3**) to increase library diversity. After the reaction, amplified barcoded DNA was purified using 0.8X AMPure XP beads (Beckman Coulter, A63881) and was eluted into a 15µL Elution Buffer. DNA concentration was measured using a Qubit™ dsDNA HS Assay (Thermo FIsher, Q32854). Final sequencing library index PCR was carried out using 20 ng of purified DNA product 2.5µL of 20µM TruR1_x_p5 primer as described in *Library indexing PCR* step. Following final library indexing, DNA was double-side purified using 0.6x-0.8x SPRIselect magnetic beads (Beckman Coulter, B23318) and eluted into 15µL of water.

### CAGE reverse-transcription reaction

Reverse-transcription reaction was set on ice and performed in PCR strip tubes. A standard 100µL reaction was composed out of 60.5µL of packed capsules, 2.5µL of 10µM RT poly(T) primer housing a 10nt UMI and 6nt barcode (**Table S3**, RT), 3µL of 10% Igepal CA-630 (Millipore Sigma, I8896-50ML), 20µL of 5X SMARTseq3 RT buffer (100mM Tris pH 8.3, 150mM NaCl, 12.5mM MgCl2, 5mM GTP, 40mM DTT), 0.5µL of 10mM dNTPs (NEB, N0447L), 2.5µL of 40U/µL Recombinant RNase Inhibitor (Takara Bio, 2313A), 1.5µL of 50µM TSO, 5µL of Maxima H-reverse transcriptase (ThermoFisher, EP0753), unless stated otherwise. Reaction volumes were scaled according to the amount of capsules used and number of reactions performed. To set up the reaction, capsules, RT primer and Igepal CA-630 (Millipore Sigma, I8896-50ML) were mixed on ice in a PCR tube, with each housing a different RT barcode. Capsules were incubated on ice for 10 min to allow the RT primer to diffuse in, followed by heating at 70°C for 5 min and cooling down on ice. Next, RT master mix, composed out of the remaining reagents was added to each tube, ensuring a thought mixing of capsules with pipetting. Tubes were incubated on ice for 5 min and the reaction was carried out at 42°C for 90 min, followed by cycling between 50°C and 42°C incubation of 2 mins for 15 cycles. Reactions were left at room temperature overnight and processed the next day.

### CAGE single-cell gDNA amplicon-Seq

K562-RFP and L1210-GFP (kindly donated by Sean McGeary and Yuyang Chen) cell suspension was mixed at a ratio of 1:1. Capsules housing single-cells were prepared as described in *Hydrogel Capsule synthesis*, except core solution contained cell suspension at a final concentration of 1.4e6 cells/mL in 1x DPBS. Resulting hydrogel capsules were resuspended in ice-cold 3x SSCI (pH 5.9). The subsequent steps were performed on ice: capsules were washed 6 times with 3x SSCI, once with TEIS (pH 7.5), three times with 3x SSCI and three times with HI10.

While the paper analyzes only gDNA amplicons, the experiment reported in the paper was simultaneously processed for joint amplicon-seq and scRNA-Seq. To this end, two 50µL reverse-transcription reactions were set on ice in two PCR strip tubes as described in *Capsule reverse-transcription reaction*.

After the RT reaction, capsules were washed three times with TI100 (pH 8.0), twice with TIbS (pH 8.0), three times with ice-cold TI100 (pH 8.0) with 2 min incubations on ice. Finally Capsules were washed twice with TI10 (pH 8.0).

The capsules containing gDNA and cDNA were then processed to amplify target gDNA regions by carrying out 16 cycles of capsule gDNA amplicon PCR. The PCR reactions were performed in 50µL volume 10µL 5X Q5U reaction buffer (NEB, M0515L), 2µL of 10mM dNTPs (NEB, N0447L), 5µL of 10µM RFP_R1 FW primer, 5µL of 10µM GFP_R1 FW primer, 5µL of 10µM universal dU CMV RV primer (**Table S3**), 20µL of packed capsules, 1µL of Q5U High-Fidelity DNA polymerase (NEB, M0515L), 2µL of 10% Igepal CA-630 (Millipore Sigma, I8896-50ML)]. The reaction was carried out by first incubating the tubes on ice for 5 mins, followed by a PCR cycling program: 98°C for 30s; 16 cycles of 98°C for 15s, 70°C for 20s, 72°C for 2 min; a final extension of 72°C for 2min and hold at 25°C. After PCR, capsules were washed once with TIbS (pH 8.0), three times with TI100 (pH 8.0) and three times with TI10 (pH 8.0). A second PCR reaction was then performed in a 50µL volume as described in Capsule dU PCR using 2.5µL of 10µM Read1_Biotin FW primer (**Table S3**), 2.5µL of 10µM universal dU CMV RV primer, 2.5µL of 10µM universal dU RV primer, 2.5µL of 10µM of TSO FW primer with 25µL of packed capsules. After the DNA amplification, capsules were washed three times with TI100 (pH 8.0), twice with TIbS (pH 8.0), three times with TI100 (pH 8.0) and twice with TI10 (pH 8.0). Washed capsules were subjected to split-and-pool barcoding as described above, with Read1_Biotin FW and TSO FW primer used in the PCR-3 barcode additions.

Following split-and-pool barcoding, capsules were split into multiple 1.5mL tubes and were dissolved as described above. DNA was purified using MinElute PCR purification kit and barcoded cDNA was eluted into 20µL of the elution buffer. Barcoded and purified DNA was amplified using PCR as described in *Post-purification barcoded DNA amplification*, using 2.5µL of TSO_fw and Read1_Biotin FW primers. After the reaction, amplified barcoded DNA was purified using 0.8X AMPure XP beads (Beckman Coulter, A63881) and was eluted into a 20µL Elution Buffer. Next, biotin pulldown using Pierce™ Streptavidin Magnetic Beads (ThermoFisher, 88816) was performed to purify amplified and barcoded biotinylated gDNA GFP/RFP amplicons. 25µL of magnetic beads were washed three times using 30µL of 5mM Tris (pH 7.4) with 0.5mM EDTA and 1M NaCl while placed on a magnetic rack. After washing, beads were resuspended in 60µL of 10mM Tris (pH 7.4) with 1mM EDTA and 2M NaCl and mixed with cDNA sample diluted to 60µL using water. Beads were incubated at room temperature for 30 min, placed on a magnetic rack. Supernatant was transferred to a new tube and was used to purify non-biotnylated DNA. Remaining beads were washed three times with 50µL of 5mM Tris (pH 7.4) with 0.5mM EDTA and 1M NaCl and biotinylated DNA was eluted by resuspending 20µL of 95% formamide with 10mM EDTA (pH 8.2) and heating ay 65°C for 5 min. Eluted DNA was purified using 1X AMPure XP beads (Beckman Coulter, A63881) and eluted into 20µL of water. Final amplicon library amplification and indexing PCR was carried using 20 μL of the purified DNA as described previously with 2.5µL of 20µM truR1_x_p5 primer that houses an 8bp library index. Following final library indexing, DNA was purified using 0.8x SPRIselect magnetic beads (Beckman Coulter, B23318) and eluted into 15µL.

### Imaging lysed cells and cDNA libraries in CAGEs

After lysis, capsules were resuspended in 3X SSCI buffer with 1X SYBR Safe DNA Gel Stain (Thermofisher, S33102). 2-5µL of packed capsules were transferred to a hemocytometer or a well of 96-well glass bottom plate and were topped with 3X SSCI buffer with 1X stain and imaged on a wide field or confocal microscopes.

### Dextran digestion reaction

Core dextran digestion reaction was performed in 1mL IMDM with 1% BSA, 0.1% F127 and 0.1% L31 pluronics unless stated otherwise in the presence of 0.5% (v/v) dextranase from *Chaetomium erraticum* (Sigma Aldrich, D0443-50ML) at 37°C for 1 min with 500 rpm shaking. Immediately after, 50µL of HFE7500 was added and capsules were spun down 300g for 1 min. All supernatant was removed and capsules were slowly resuspended in 500µL of media and transferred to a new tube without disturbing the bottom HFE7500 phase. Capsules were washed once with 1mL IMDM with 1% BSA, 0.1% F127 and 0.1% L31 pluronics.

### CAGE single cell RNA-Seq

In the optimized protocol, Capsules housing single-cells were prepared as described in *Hydrogel Capsule synthesis*, except the core solution was prepared in DPBS and contained 8mM DTT, 1% BSA and a final concentration of 1.4e6 cells/mL. Resulting hydrogel capsule pellet was resuspended in room temperature 1mL IMDM with 1% BSA, 0.1% F127 and 0.1% L31 pluronics and was washed twice with 300g spins for 1 min. Next, core dextran was removed as described in *Dextran digestion reaction*. Following this, all steps were performed on ice. First, capsules were washed three times with an ice-cold 3X SSCI buffer with 800g spins for 1 min. Cells were lysed by washing with ice-cold 7.5pH TEIS with 5 min incubations on ice. After lysis, capsules were imaged (check above), washed three times with 3X SSCI and three times with HI10, packed at 800g for 1 min and stored on ice.

Reverse-transcription reaction was set on ice and performed in PCR strip tubes as described in *Capsule reverse-transcription reaction*. After the RT reaction, capsules were washed three times with TI100 (pH 8.0), twice with TIbS (pH 8.0), three times with ice-cold TI100 (pH 8.0) with 2 min incubations on ice. Finally capsules were washed twice with TI10 (pH 8.0).

Next, Capsule dU PCR was performed to carry out cDNA amplification as described previously, using, 5µL of 10µM TSO FW primer, 5µL of 10µM dU universal RV primer (**Table S3**), After the cDNA amplification, capsules were washed three times with TI100 (pH 8.0), twice with TIbS (pH 8.0), three times with TI100 (pH 8.0), twice with TI10 (pH 8.0). Capsules were imaged on a confocal microscope to inspect the cDNA signal (from labeled dU primer). Washed capsules are subjected to split-and-pool barcoding as described above with TSO FW primer used in the PCR-3 barcode additions.

Following split-and-pool barcoding, capsules were split into multiple 1.5mL tubes and were dissolved as described above. DNA was purified using MinElute PCR purification kit and barcoded cDNA was eluted into 20µL of the elution buffer. Barcoded and purified DNA was amplified using PCR as described in *Post-purification barcoded DNA amplification*, using 2.5µL of TSO FW primer. After the reaction, amplified barcoded cDNA was purified using 0.8X AMPure XP beads (Beckman Coulter, A63881) and was eluted into a 15µL Elution Buffer. DNA concentration was measured using QuBit fluorometer and cDNA fragment length profile was assessed using Agilent Tapestation D5000 kits (Agilent, 5067).

To construct Illumina sequencing libraries, 50 ng of barcoded and amplified cDNA was fragmented, blunted and dA-tailed by mixing it with 3.5 μL of NEBNext Ultra II FS Reaction buffer (NEB), 1 μL of fully mixed NEBNext Ultra II FS Enzyme mix (NEB) and nuclease free water to a final volume of 17.5 μL. The reaction mix was vortexed for 10 s, spanned down and placed into a preheated PCR machine for incubation at 37°C for 10 min; 65°C for 30 min; hold 4°C (lid at 75°C). Fragmented DNA was purified using 0.8X AMPure XP beads (Beckman Coulter, A63881) and eluted into 16µL of water. Single-stranded DNA adapter, containing Illumina’s TruSeq read 1 (Adp. TruSeq R1 FW and RV) sequences were annealed together at the final concentration of 1.5 μM by mixing 1.5 μL of forward ligation primer and reverse ligation primer (each at 100 μM) with 97 μL of nuclease-free water and heating the solution to 95°C for 2 min and cooling it at room temperature for 5 min. 16 μL of the purified fragmented DNA was mixed with 1.25 μL of 1.5 μM annealed adapters, 1.5µL of 10X T4 DNA ligase buffer, 15 μL of NEBNext Ultra II Ligation Master Mix (NEB, E7805S kit) and 0.5 μL of NEBNext Ultra II Ligation Master Enhancer (NEB, E7805S kit). The ligation reaction was carried out by incubating in the thermocycler for 15 min at 20°C (lid at 30°C). After ligation, the reaction mix was diluted to 100 μL and was purified using AMPure XP beads (Beckman Coulter, A63881) with 0.8X volume ratio and eluted into 40 μL nuclease-free water.

Final sequencing library indexing PCR was carried out as described previously with 2.5µL of 20µM truR1_x_p5 primer that houses an 8bp library index.

Following final library indexing, DNA was double-side purified using 0.6x-0.8x SPRIselect (Beckman Coulter, B23318) magnetic beads and eluted into 15µL

### CAGE ATAC-Seq

While the paper analyzes only the accessible gDNA amplicons, the experiment reported in the paper was simultaneously processed for joint ATAC-seq and RNA-Seq in single cells. To this end, a K562, L929 (kindly donated by Sean McGeary) and L1210 cell suspension was prepared by mixing cells at a ratio of 1:1:1. Capsules housing single-cells were prepared as described in *Hydrogel Capsule synthesis*, except the core solution contained 8mM DTT, 1% BSA and final concentration of 1.4e6 cells/mL in 1x IMDM. After 45 photopolymerization hydrogel capsules were purified as described previously and the resulting hydrogel capsule pellet was resuspended in room temperature 1mL IMDM with 1% BSA, 0.1% F127 and 0.1% L31 and washed twice with 300g spins for 1 min. Next, core dextran was removed as described in *Dextran digestion reaction*.. Following this, all steps were performed on ice.

Unloaded Tn5 transposase (Diagenode, C01070010-20) was loaded with adapter sequences carrying a barcode (**Table S3**, Tn5). First, the oligos were resuspended to 100µM in 40mM Tris (pH 8.0) with 50mM NaCl, universal Tn5_rev oligo was mixed with oligo Tn5_B0_x or Tn5_oligoB at 1:1 ratio and annealed by in a thermocycler: 95°C, 5min; cool to 65°C (-0.1°C/s); 65°C, 5min; cool to 4°C (-0.1°C/s). Annealed adapters were mixed at ratio 1:1 and 10µL were gently added to 10µL of unloaded Tn5. The solution was incubated at 23°C for 30 min in a thermocycler and 10µl of 99% glycerol was added to enable the sorting of loaded Tn5 at -20°C.

To perform gDNA tagmentation, 21.6µL of packed capsules were mixed with 30µL of 2x TD buffer (20 mM Tris 10 mM MgCl2; 20% (vol/vol) DMF, pH 7.6), 2.4µL of loaded Tn5 and 6µL of 10X Lysis additive (5% Tween20, 0.05% Digitonin in DPBS). The solution was incubated on ice for 5 min, followed by incubation at 37°C for 30 min. After the reaction, capsules were washed three times with TEIS (pH 7.5) and then Proteinase K treated in 200µL of TEIS (pH 7.5) with 10µL of Proteinase K (NEB, P8107S) for 15 min at 55°C. After the reaction, capsules were washed three times with TI100 (pH 7.5) and twice with TI (pH 7.5).

Next, Reverse-transcription reaction was set on ice and performed in PCR strip tubes as described in *Capsule reverse-transcription reaction*. After the RT reaction, capsules were washed three times with TI100 (pH 8.0), twice with TIbS (pH 8.0), three times with ice-cold TI100 (pH 8.0) with 2 min incubations on ice. Finally capsules were washed twice with TI10 (pH 8.0).

Next, Capsule dU PCR was performed as described before, using 2.5µL of 20µM NexteraR1_x_p5, 2.5µL of 20µM TSO FW (**Table S3**), 5µL of 10µM dU universal RV primer with 25µL of packed capsules. After the PCR, capsules were washed twice with TI100 (pH 8.0), twice with TIbS (pH 8.0) and twice with TI10 (pH 8.0). Resulting tagmented and amplified DNA was barcoded by using the split-and-pool approach as described previously with P5 Biotin FW and TSO FW primer used in the PCR-3 barcode additions.

Following split-and-pool barcoding, capsules were dissolved in 1.5mL tubes as described above. DNA was purified using MinElute PCR purification kit and barcoded DNA was eluted into 20µL of the elution buffer. Barcoded and purified DNA was amplified using PCR as described in *Post-purification barcoded DNA amplification*, using 1.25µL of 20µM P5 Biotin FW and 1.25µL of 20µM TSO FW primer. DNA was purified using AMPure XP beads (Beckman Coulter, A63881) with 1.8X ratio and eluted into 20µL. Next, biotin pulldown using Pierce™ Streptavidin Magnetic Beads (ThermoFisher, 88816) was performed to purify amplified and barcoded biotinylated gDNA fragments from non-biotinylated cDNA libraries as described previously.

Next, indexing with qPCR quantification was run using 25µL 2x NEBNext® High-Fidelity 2X PCR Master Mix (NEB, M0543L), 2.5µL of 20µM P7_rv primer, 2.5µL of 20µM p5 FW primer and 20µL of purified barcoded DNA. Reactions were performed using a PCR cycling program: 98°C for 50s; 6 cycles of 98°C for 10s, 68°C for 20s, 72°C for 1min; a final extension of 72°C for 4min and hold at 4°C and the reaction was transferred on ice. Next, 5µL of partially amplified DNA was mixed with 0.25µL of 20µM of each primer, 0.1µL of 100X SYBRgreen, 5µL of 2x NEBNext® High-Fidelity 2X PCR Master Mix (NEB, M0543L) and 4.4µL of water. The reaction was carried out on a qPCR machine using the same PCR program as for the initial 6 cycles, except imaging the samples after every extension step. RFU vs cycle plot was used to determine the number of cycles needed to reach ⅓ of maximum intensity. This number of cycles was used to amplify the remaining 45µL of partially amplified DNA. After the reaction, DNA was purified using double sided size AMPure XP (Beckman Coulter, A63881) size selection with a bead ratio of 0.5X-1.3X. Final library DNA was eluted to 15µL of EB, quantified using Qubit and qPCR before sequencing

### Fixation of cells in suspension

Cell suspensions were centrifuged at 400g for 3 mins and the pellet was resuspended in 1mL of 4% PFA in DPBS. The solution was incubated at room temperature for 15 mins, followed by 3 washes with 1x DPBS + 0.1% BSA. The fixed cells were encapsulated into capsules by adding the cells to ∼1.4e6 cells/mL into the core solution.

### Reverse cross-linking cells in capsules

Fixed cells were reverse cross-linked in capsules by washing the capsules twice with 3x SSCI + 0.01U/µL RiboLock RNAse Inhibitor (ThermoFisher, EO0381). Next, Capsules were washed three times with 100mM Tris (pH 7.5) with 10mM EDTA, 0.1% Igepal CA-630 (Millipore Sigma, I8896-50ML), 0.1% SDS. Finally, capsules were resuspended in the same solution to a volume of 1 mL and 50 µL of proteinase K (NEB, P8107S) was added followed by incubation at 56°C for 30 min. After the reverse crosslinking, capsules were washed three times with 3X SSCI, three times with HI10 and were used in a reverse transcription reaction as described above (*Capsule single-cell RNA-seq*).

### Freezing capsules

Capsules were frozen at -80°C in the presence of 10% DMSO. Capsules containing cell lysates were frozen in a 3X SSCI buffer with 0.05U/µL of RNAse inhibitor (NEB, M0314S). Capsules containing DNA only were frozen in TI (pH 8.0) with 10% DMSO.

### Preparation of PBMCs

Human Peripheral Blood Mononuclear Cells were obtained from Lonza (CC-2704). The cryo-preserved cell vials were stored in liquid Nitrogen until use, and then thawed in a 37°C metal bead bath for 7 minutes. The thawed cell suspension was transferred into a 15 mL falcon tube. 1 mL of 37°C pre-warmed DMEM (ThermoFisher, 10566016) with 10% FBS (ATCC, 30-2020) and 1x Penicillin/Streptomycin (ThermoFisher, 10378016) was slowly added into the original cryopreservation tube. This media was then added one drop at a time into the falcon tube. Additional 8 mL of pre-warmed media were added slowly onto the cell suspension. Cells were resuspended gently and centrifuged at room temperature at 300g for 4 mins. All of the media was aspirated from the cell pellet and cells were suspended in 5 mL of fresh, 37°C appropriate media with 10% FBS and 1x Penicillin/Streptomycin and transferred into 25 cm^2^ culture flask with a filter cap (Corning). Cells were cultured in a 37°C incubator with 5% CO2. In the case of PBMCs, cells were washed one more time with IMDM + 10% FPBS +1x Pen/Strep and were passed through a 40µm strainer. Following this, cells were washed with ice cold plain IMDM with 0.5% BSA. PBMC were encapsulated at a final concentration of 1.4e6 cells/mL in the core mix and then processed for scRNA-Seq as described above.

### Mammalian cell line culture

K562, L1210 and L929, Jurkat cell lines were cultured and used in experiments described in this work. Unless otherwise stated, K562 and L1210 cells were cultured in IMDM (Thermo Fisher, 10566016) + 10% FBS (ATCC, 30-2020) + 1x penicillin/streptomycin (Thermo Fisher, 10378016) in T25 flasks (Corning). L929 cells were cultured in EMEM (ATCC, 30-2003) + 10% horse serum (Thermo Fisher, A5669502) + 1x penicillin/streptomycin (Thermo Fisher, 10378016) in T25 flasks (Corning). Jurkat and HEK293T cells were cultured in RPMI1640 (Thermo Fisher, 61870036) + 10% FBS (ATCC, 30-2020) + 1x penicillin/streptomycin (Thermo Fisher, 10378016) in T25 flasks (Corning). Cell lines were cultured until they reached ∼80-90% confluency. For passaging, suspension cells (K562, L1210, Jurkat) were resuspended gently and collected into a 15 mL tube. 5 mL of 37°C pre-warmed complete media was used to wash the flask and was added into the same 15 mL tube. Adherent cells (HEK293T, L929) cells were passaged by aspirating the used media and washing 2 times with 2 mL of 37°C pre-warmed 1x DPBS buffer. Cells were detached by adding 1 mL of TrypLE reagent (Thermofisher Scientific, 12605010), waiting 10s and quickly aspirating all of the TrypLE. Cells were incubated in 37°C for 2 minutes and were resuspended in 5 mL of 37°C pre-warmed complete media. Cells were centrifuged at 300g for 4 min, followed by media aspiration and resuspension in 5 mL of fresh pre-warmed complete media. Cells were counted using an automatic cell counter and 500,000 cells were seeded into 5 mL of 37°C pre-warmed complete media in a 25cm2 flask. For encapsulation experiments, cells were washed twice with ice cold DPBS with 0.1% BSA or room temperature IMDM with 0.1% BSA and were encapsulated at a final concentration of 1.4e6 cells/mL in core mix and then processed immediately for scDNA-seq/scRNA-Seq/scATAC-seq as described previously or grown into colonies as described below.

### iPSC culturing

Induced pluripotent cells were cultured in a 6-well plate on a layer of Matrigel (Corning) in mTeSR media (STEMCELL Technologies, 100-0276). To passage the culture, cells were detached by removing media, rinsing twice with DPBS, adding 1mL of Accutase (STEMCELL Technologies, 07920) and incubating at 37°C for 7 min. Next, cells were resuspended in 5mL of plain DMEM and centrifuged in a swinging bucket centrifuge for 4 min at 300g. Supernatant was aspirated and the cell pellet was resuspended in 1mL of DMEM. 2mL of Matrigel in DMEM solution was added to a single 6-well plate well. The plate was incubated in 37°C for 30 min. Any remaining solution was aspirated and 500K of iPSC cells in DMEM were seeded into 2mL of mTeSR (STEMCELL Technologies, 100-0276) with 10µg/mL Y-27632 (Dihydrochloride) (STEMCELL Technologies, 72302). After a day of culturing, media was changed with fresh mTeSR without Y-27632. Media changes were performed daily for 3-4 days, till cells reached 80-90% confluency.

### Cell-line culturing in CAGEs

Capsules housing single cells were prepared as described in *Hydrogel Capsule synthesis*, except the core solution was prepared in 1x IMDM and contained 8mM DTT, 1% BSA, and a final concentration of 1.4e6 cells/mL. Emulsion was polymerized for 45s and capsules were purified as described previously. Resulting hydrogel capsule pellet was resuspended in 37°C temperature 1mL IMDM with 10% FBS, 0.1% F127 and 0.1% L31 pluronics, washed twice with 300g spins for 1 min and core dextran was removed as described previously. Capsules were washed once with 1mL IMDM with 10% FBS, 0.1% F127 and 0.1% L31 pluronics. Packed capsules were transferred to wells of 24-well plate and cultured in 1mL of complete media as described in mammalian cell line culturing part above.

To change media, capsules were resuspended in the growth medium and were transferred to a reverstable 37µm strainer (STEMCELL technologies, 27215). Capsules were washed in the strainer with 3mL of complete growth medium. Next, the strainer was inverted and the capsules were eluted back into a well of a 24-well plate using 1mL of complete growth medium. Any manipulation of capsules was done in complete growth media with 0.1% F127 and 0.1% L31.

### iPSC culturing in CAGEs

Capsules housing single iPSCs were generated using the protocol described above with a final concentration of 1.4e6 cells/mL in the core mix. Capsule clean-up and dextran removal were done as described above. After clean-up, capsules were washed twice with mTeSR 3D + 0.1% F127, 0.1% L31 + 0.1% BSA. Packed capsules were transferred to wells of 24-well plate and cultured in 1mL of mTeSR 3D media (STEMCELL Technologies, #03950) with 1X CloneR^TM^2 cloning supplement (STEMCELL Technologies, 100-0691). To change media, capsules were resuspended in the surrounding growth medium and were transferred to a reverstable 37µm strainer (STEMCELL Technologies, 27215). Capsules were washed in the strainer with 3mL of IMDM with 0.1% F127 + 0.1% L31. Next, the strainer was inverted and the capsules were eluted back into a well of a 24-well plate using 1mL of mTeSR 3D media. Any manipulation of capsules was done in complete growth media with 0.1% F127 and 0.1% L31.

### mHPC culturing in capsules

LSK mHPCs were gifted by the lab of Fernando Camargo (Harvard). Capsules housing single mHPCs were generated using the protocol described above with a final concentration of 0.8e6 cells/mL in the core mix and no BSA. Capsule clean-up and dextran removal were done as described above, except all washes were performed in plain IMDM + 0.1% F127 and 0.1% L31. After clean-up, capsules were washed twice F12 media (Thermo Fisher, 11765054) supplemented with 10mM HEPES, 1X P/S/G, 1 mg/ml PVA, 100 ng/ml TPO (ThermoFisher, 300-18-10UG), 10 ng/ml SCF (Thermo Fisher, 300-07-10UG), 1x ITSX (Thermo Fisher, 51500056), 0.1% F127, 0.1% L31. Packed capsules were transferred to wells of 24-well plate and cultured in 1mL F12 expansion media. Media was exchanged every two days by collecting all the capsules into a 1.5mL tube, spinning down at 300g for 1 min and resuspending in fresh media.

### Fixing and staining clones in CAGEs

Colonies in capsules were fixed by resuspending packed capsules in 4% PFA solution in DPBS and incubating at room temperature for 20 min. Capsules were washed twice and stored in ice-cold DPBS with 0.1% BSA, 0.1% F127, 0.1% L31. Cells were permeabilized by adding Triton X-100 to a final concentration of 0.1% and incubating for 10 min. Capsules were washed twice and stored in a fridge. For iPSC colonies, immunostaining was performed against SOX2 by adding 2µL of Alexa Fluor® 488 anti-SOX2 Antibody (BioLegend, 656109) (0.5mg/mL) to 600µL of capsules solution in DPBS with 0.1% B/F/L. As a control, 2µL of 0.5mg/mL FITC anti-mouse CD45.1 Antibody (BioLegend, 110705) was used. Capsules were stained on ice for 15 min, followed by 3 washes with DPBS + 0.1% BSA/F127/L31 with 5 min incubation on ice between washes. Stained capsules were imaged on a confocal microscope.

### Treatment with HDAC and DNMT1 inhibitors

Survival curves for DNMT inhibitors [Decitabine (Dec) and 5-aza-cytidine (Aza)], and the HDAC inhibitor Vorinostat (Vor) (MedChemExpress, HY-A0004, HY-10586, HY-10221) were generated by seeding 15,000 (L1210) and 20,000 (K562) into 96-well plates and growing the cells in 100µL of IMDM + 10% FBS, 1X Pen/Strep in the presence of 0, 0.001, 0.003, 0.01, 0.03, 0.1, 0.3, 1, 3, 10 µM of each drug. Cells were passaged every two days by transferring 15µL of cells into a fresh 85µL of media loaded with the same concentration of drugs. Cell viability and number was evaluated every two days using a confocal microscope by mixing 7.5µL of cell culture with 42.5µL of DPBs with 1X DAPI stain.

### CAGE clonal expansion and inc-RNA-seq

Cells were pre-treated with drugs by seeding 150,000 (L1210) and 200,000 (K562) cells into 6-well plate wells (Corning). Cells were grown in 2mL of IMDM with 10% FBS, 1X Pen/Strep in the presence of 0.05 µM Dec, 0.7µM Aza, 0.4µM Vor (MedChemExpress) or no drugs for 48 hours. Next, cells were collected and K562 and L1210 cell-lines were pooled based on the drug condition at a 1:1 ratio. Cell suspension was encapsulated into hydrogel capsules as described above. Each drug condition was encapsulated for 2 hours, collecting the droplets into four 30 min fractions. Hydrogel capsules were polymerized, washed and dextranase treated as described above. One fraction of each condition was sampled instantly by lysing the cells inside the capsules - capsules were washed three times with ice-cold 3X SSCI buffer with 800g spins for 1 min. Cells were lysed by washing twice with ice-cold TEIS (7.5pH) with 5 min incubations on ice. After cell lysis, capsules were frozen down at -80°C in 3X SSCI with 10% DMSO and 0.05U/µL Murine RNAse Inhibitor (NEB, M0314S). Remaining fractions of packed hydrogel capsules were seeded into a different 24-well plate well into 1mL of IMDM with 10% FBS and 1x Pen/Strep, 0.1% F127 and 0.1% L31 with appropriate drug. 10,000 K562 and L1210 cells from appropriate drug conditions were seeded outside the capsules for the first two days. Every two days, media was exchanged by collecting the capsules using an reversable a 40µm strainer (STEMCELL Technologies) as described above and eluting into fresh IMDM with 10% FBS and 1x Pen/Strep, 0.1% F127 and 0.1% L31 with appreciate drug. Every 2 days, one well of each condition was sampled for Capsule RNA-seq by lysing the cells and freezing at -80°C as described above. After all the time points were collected, hydrogel capsules were thawed, washed once with 3X SSCI, once with TI10 (pH 7.5), once with TI (pH 7.5). Next, capsules were treated with Turbo DNAse I (Thermo Fisher) by adding 8µL of 2U/µL enzyme to 300µL of capsules in a 1X reaction buffer and incubating at 37°C for 10 min. Capsules were then washed once with 3X SSCI and TEIS (pH 7.5) and resuspended in 500µL of TEIS (pH 7.5). Next, 8µL of Proteinase K (NEB, P8107S) were added to each tube, followed by 15 min incubation at 37°C. Finally, capsules were washed three times with 3X SSCI buffer and three times with HI10. Resulting capsules were used in a capsule RNA-seq protocol as described above. Resulting barcoded cDNA libraries were purified, fragmented and indexed with 48 indices as described above.

### Sequencing

DNA library fragment size was assessed using Agilent Bioanalyzer hsDNA or Tapsestation D5000 kits (Agilent). Sequencing libraries, housing different P5 indices were pooled, and concentration was assessed using a qPCR using KAPA SYBR® FAST Universal qPCR Kit (KAPA Biosystems) according to manufacturer’s guidelines. All libraries were sequenced on a Paired-end Illumina Next-Generation sequencing platform. See **Table S4** for sequencing kit and parameters

## DATA ANALYSIS

### Software packages

Except as stated otherwise below, all data analysis was carried out in Python (3.10) with the packages scanpy (1.10.4), numpy (1.26.4), pytorch (2.2.2), pymc (5.20.0), scikit-learn (1.6.1).

### Sequencing read demultiplexing and pre-processing

BCL sequencing files received from Illumina BaseSpace portal were converted to raw FASTQ files for each of the 4 (R1, R2, R3, R4) sequencing reads without read trimming or masking by using bcl2fastq software (Illumina, v2.20.0.422). Raw FASTQ files were processed using a custom pipeline adapted from inDrops.py (Zilionis et al., 2017). The new code, inCaps.py is available at github.com/AllonKleinLab/paper-data/tree/master/Mazelis2025_CAGEs.

In brief, sequencing reads were demultiplexed based on Read1 index sequence, and were filtered based on expected structure (all barcodes matching a barcode whitelist with hamming distance of 1). The demultiplexed reads were then processed using Trimmomatic (*1*) (with parameters: LEADING 28; SLIDINGWINDOW 4:20; MINLEN 20), followed by Poly-A length trim to 4 nt and selecting reads that are longer than MINLEN. For surviving reads, the fraction of length composed out of same base repeats (only considering > 5) was determined, rejecting reads whose fraction is greater than 0.5. As output, for every input run part (sequencing lane), this produces a filtered FASTQ file for every library index contained in that run. Next, all filtering results were combined to identify abundant cell barcodes and create a summary filtering table for each library processed. An index is created to sort the reads according to the name of their barcode of origin. Barcodes with less than 250 total reads (across all library-run-parts) are ignored, and placed at the end of the file, creating gzipped and sorted FASTQ files with headers containing capsules barcodes (and UMI if applicable) and an index of the byte offsets for every barcode with more than 250 reads. Demultiplexed, filtered and sorted FASTQ files were used for mapping and counting as described in the following sections.

### Read mapping for amplicon sequencing

Sequencing reads were first processed as described above to generate demultiplexed, filtered and sorted FASTQ files. Sequencing reads were aligned to a GFP or RFP reference using STARSolo (*2*) counting the number of reads aligning to each reference for each detected barcode.

### Capsule RNA-Seq count matrix generation

Sequencing reads were first processed as described above to generate demultiplexed, filtered and sorted FASTQ files. Count matrices were generated using STARsolo pipeline (*2*) (STAR 2.7.9a) with the with parameters (*--soloFeatures Gene, GeneFull, --genomeSAsparseD 3, --outSAMtype BAM SortedByCoordinate, --outSAMattributes CB CR UR UB UY GN GX, -- outFilterScoreMinOverLread 0.3, --outFilterMatchNminOverLread 0.3, --soloCBmatchWLtype 1MM_multi_Nbase_pseudocounts, --soloUMIfiltering MultiGeneUMI_CR, --soloUMIdedup 1MM_CR, --soloMultiMappers EM, --soloCellFilter EmptyDrops_CR, --quantMode TranscriptomeSAM GeneCounts*) or InDrops pipeline with Bowtie 1.2.2 (m = 200; n = 1; l = 30; e = 70) unless stated otherwise. Human GRCh38.101 and mouse GRCm.101 genome assemblies were used.

### Cell-line Capsule scRNA-seq of data analysis

Count matrices were generated as described above using the STARsolo pipeline. Count matrices were filtered by selecting barcodes with >3000 UMIs and fraction of mitochondrial counts being below 0.2. Counts were quantile-quantile (Q-Q) downsampled among all samples. Gene-expression correlation analysis was performed after transcript per million (TPM) normalization and log10(1+TPM) scaling.

### PBMC Capsule scRNA-seq data analysis

Count matrices were generated using the STARsolo pipeline with intronic reads as described above. Count matrices were filtered by selecting barcodes with >2000 UMIs and fraction of mitochondrial counts being below 0.2. Counts were normalized to CP10K, and z-scored. Scanpy was used to identify high variable genes (sc.pp.highly_variable_genes with flavor=’seurat’, min_mean=0.0125, max_mean=3, min_disp=0.2) and to perform PCA, UMAP and Leiden clustering.

### Data comparison to 10x Genomics

Capsule scRNA-seq HEK293t, NIH 3t3 and PBMC data was compared to an equivalent sample of 10X Genomics scRNA-seq data downloaded from the companies website in raw FASTQ format. Count matrices for Capsule and 10X Genomics data were generated using STARsolo pipeline with the same parameters (*--soloType CB_UMI_Simple, --soloUMIlen 12, --soloFeatures Gene, -- genomeSAsparseD 3, --outSAMtype BAM SortedByCoordinate, --outSAMattributes CB CR UR UB UY GN GX, --outFilterScoreMinOverLread 0.3, --outFilterMatchNminOverLread 0.3, -- soloCBmatchWLtype 1MM_multi_Nbase_pseudocounts, --soloUMIfiltering MultiGeneUMI_CR, -- soloUMIdedup 1MM_CR, --soloMultiMappers EM, --soloCellFilter EmptyDrops_CR, --quantMode TranscriptomeSAM GeneCounts*). Human GRCh38.101 and mouse GRCm.101 assemblies were used. Count matrices were filtered by selecting barcodes with >3000 UMIs and fraction of mitochondrial counts being below 0.2. Total counts were quantile-quantile (Q-Q) downsampled among all samples. Gene-expression correlation analysis was performed after transcript per million (TPM) normalization and log10(1+TPM) scaling. To compare gene-expression for different transcript length, *gffutils* was used to extract transcripts length from GTF files used to make the genome reference. Gene expression was compared between platforms at different length cutoffs. To generate UMI saturation curves, random sampling was used to sample barcode-umi-gene combinations from output aligned and sorted bam files to generate a table of number of UMIs and reads for each barcode at different sequencing read depths.

### Capsule ATAC-Seq data processing

Capsule ATAC-seq approach used pair-end sequencing to map out gDNA fragment length. While Read1 sequencing reads contain only the gDNA fragment, Read4 contains capsule barcodes followed by a gDNA fragment. First, parts of Read4 sequencing reads corresponding to gDNA fragments were extracted using *seqtk trimfq* command with a parameter *-b 50*. Sequencing reads were then processed as described above to generate demultiplexed, filtered and sorted FASTQ files twice, once for each gDNA read (Read1 and Read4). Next, *fastq-pair* was used to pair the two demultiplexed and filtered gDNA fragment FASTQ files, which were then merged using *NGmerge*. Merged FASTQ files were aligned to merged human GRCh38.101 and mouse GRCm.101 index using *bowtie2* (with parameter --very-sensitive, -p 4) and bam files were sorted using *samtools sort* function. Mitochondrial reads were counted and removed, followed by BAM file deduplication, sorting and indexing using *samtools*. Next a pysam script was used to move barcode sequences from BAM read headers to bam CB tag and a string tag was appended to each barcode to label runs corresponding to different indices. Following this, bam files were merged for all the runs and *Macs3* was used to call peaks (*--nomodel, --shift “-100”, --extsize “200”, -B, --SPMR, --keep-dup “all”*). *HTSeq* and *SnapATAC* python package was used to analyze the processed Capsule ATAC-seq data.

### Capsule colony RNA sequencing

#### Preprocessing

Read mapping and cell filtering based on counts and mitochondrial fraction were carried out as described above. Raw counts were then normalized to 10,000 counts per cell (CP10K). Highly variable genes were selected using Scanpy’s implementation of the Seurat v3 approach (sc.pp.highly_variable_genes with flavor=’seurat_v3’; 4,000 genes for K562 cells and 5,000 genes for L1210 cells; using unnormalized counts for variance calculations).

#### Mock colony generation

Mock-colony size-matched controls were generated using a count-based resampling approach. Let *n*_1_ …, *n_M_*} be the total counts for the *M* cells sampled for a given timepoint and condition in the real data. For each value of *n_k_*, we generated a mock cell with the same total counts by (1) randomly sampling single cells (time point 0 transcriptomes) from the corresponding condition, (2) combining raw unnormalized counts to match or exceed target colony sizes, with a minimum of 2*^t^*^/days^ cells from time point 0 combined for mock colonies at time *t*, and enough cells such that that the total counts equal or exceed *n_k_*; then (3) downsampling the counts to *n_k_*. The mock colonies were then processed as the real colonies, except where stated otherwise below.

#### Visualizing variable genes in colonies

Gene coefficient of variation (CV) vs mean plots were generated by first calling scanpy’s sc.pp.highly_variable_genes as described above to obtain means, variance and normalized variances for every time point and replicate in for control (untreated samples), for real data and mock clones. The values for each gene were then weighted averaged over the three control replicate samples for colonies from days 4 and 6, weighted by the number of colonies sampled per library. Gene-gene correlation maps were similarly generated for the real data by first calculating pairwise gene correlations within each sample, and then obtaining the weighted average correlation across the three untreated replicates. The top 150 variable genes were selected after filtering out mitochondrial, ribosomal and other predefined gene sets using exclusion lists. For the heatmap of the mean correlations, the genes were hierarchically clustered using average linkage clustering.

#### Dimensionality reduction

Non-negative matrix factorization (NMF) was performed on the highly variable genes using scikit-learn’s NMF implementation with the following parameters: init=’nndsvd’, random_state=0, n_iter=400, alpha_W=0, alpha_H=0, l1_ratio=1.0, and tol=1e-3. The solver was set to ‘cd’ (coordinate descent). NMF was carried out on log10(1+CP10k)-transformed counts. To determine the optimal number of factors, we compared explained variance ratios between real and permuted data across 1-20 components (*3*), with 5 permutations and sampling 10% of cells for computational efficiency. This procedure identified the top 15 components as explaining variance above-random and this number of NMF programs was used for both cell lines.

#### Visualization by UMAP embedding

For visualization only, batch effects from library preparation and cell seeding were corrected using Harmony (Korsunsky et al., 2019) using the scanpy implementation, with the following non-default parameters: theta=4.0, lambda=10.0, and sigma=0.05. Harmony was applied to the 15 NMF program usages per cell. A nearest neighbor graph was constructed using the Harmony-corrected program usages with n_neighbors=10, and scanpy’s UMAP was then applied to the batch-corrected embedding with default parameters.

#### Heatmaps of NMF gene loadings

For heatmaps of gene contributions to NMF programs, the NMF loading matrix (“H” matrix in scikit-learn) was normalized by dividing each gene’s loading by its mean non-zero expression level (calculated as mean expression when expressed / fraction of cells expressing). The genes shown in the final heatmaps were filtered to remove mitochondrial genes and non-coding genes to improve interpretability, and ordered by descending normalized loading values.

#### NMF usage means, CVs and Fano Factors

NMF program usage values were first normalized for each cell such that the sum of usages across all programs equaled 1. The mean, CV, and Fano factor of each program’s usage across colonies was then calculated at each timepoint. To calculate mock colony CV and Fano factors, we corrected for systematic differences in means over time by regressing the CVs using the power law relationship observed between mean and CV. For comparisons across conditions, each metric was normalized to its value in control colonies at the same timepoint.

#### Machine learning program ON/OFF switching rates

A model of stochastic state transitions within cells of an expanding clone was fit to the distribution of usages for each NMF program over time, in each condition. Referring to Supplemental Text for details on the model, the parameters are: the on rate (*r*_01_/ cell cycle), the off rate (*r*_10_/ cell cycle), the initial state probability (*p*_0_), the NMF program usage background level (ε), and the NMF program usage expression scale and noise parameters (*a, b*, σ, σ_0_). These parameters were inferred by Markov Chain Monte Carlo (MCMC) sampling using PyMC, with a prior distribution and likelihood as defined in Supplemental Text 1. To generate the likelihood, we carried out 1.6 ⋅ 10^6^ simulations of stochastic cell state switching with different rates (*r*_01_, *r*_10_) and then trained a surrogate neural network model (2 hidden layers: 64, 32 nodes) to predict the number of cells in states (0,1) in each clone over time *t*, enabling rapid likelihood computation during MCMC inference. Parameters were inferred using 8 parallel MCMC chains of 2,000 samples each after 2,000 tuning steps. Convergence was assessed using the Gelman-Rubin statistic. Rates (*r*_01_,*r*_10_) increasing following drug treatment imply shorter persistence time for a given NMF program, while the opposite implies longer persistence. Code implementing simulations, surrogate model training and MCMC inference will be available at github.com/AllonKleinLab/paper-data/tree/master/Mazelis2025_CAGEs.

